# BamQuery: a proteogenomic tool for the genome-wide exploration of the immunopeptidome

**DOI:** 10.1101/2022.10.07.510944

**Authors:** Maria Virginia Ruiz Cuevas, Marie-Pierre Hardy, Jean-David Larouche, Anca Apavaloaei, Eralda Kina, Krystel Vincent, Patrick Gendron, Jean-Philippe Laverdure, Chantal Durette, Pierre Thibault, Sébastien Lemieux, Claude Perreault, Grégory Ehx

## Abstract

MHC-I-associated peptides (MAPs) derive from selective yet highly diverse genomic regions, including allegedly non-protein-coding sequences, such as endogenous retroelements (EREs). Quantifying canonical (exonic) and non-canonical MAPs-encoding RNA expression in malignant and benign cells is critical for identifying tumor antigens (TAs) but represents a challenge for immunologists. We present BamQuery, a computational tool attributing an exhaustive RNA expression to MAPs of any origin (exon, intron, UTR, intergenic) from bulk and single-cell RNA-sequencing data. We show that non-canonical MAPs (including TAs) can derive from multiple different genomic regions (up to 35,343 for EREs), abundantly expressed in normal tissues. We also show that supposedly tumor-specific mutated MAPs, viral MAPs, and MAPs derived from proteasomal splicing can arise from different unmutated non-canonical genomic regions. The genome-wide approach of BamQuery allows comprehensive mapping of all MAPs in healthy and cancer tissues. BamQuery can also help predict MAP immunogenicity and identify safe and actionable TAs.

## 1. INTRODUCTION

The immunopeptidome is the repertoire of MHC-I-associated peptides (MAPs) that represents in real-time the landscape of the intracellular proteome as it is molded by protein translation and degradation^1^. In recent years, immunopeptidomic data has been harvested to identify relevant and targetable tumor antigens. Indeed, MAPs deriving from mutations characterizing the neoplastic transformation (mutated tumor antigens (TA), also known as neoantigens) can be recognized by cytotoxic T cells and used as anti-cancer therapeutic targets^2^.

The immunopeptidome is typically assumed to result from the degradation of canonical proteins, coded by exons and translated from known open-reading frames. However, recent proteogenomics (proteomic informed by genomics such as RNA sequencing (RNA-seq)) findings evidenced that ∼5-10% MAPs can also derive from non-canonical (nc) regions of the genome, such as introns, non-coding RNAs (ncRNA) or endogenous retroelements (EREs), as well as from out-of-frame translation of exons^3-5^. While 99% of somatic mutations are located in non-coding regions^6^, the vast majority of the discovered ncMAPs are non-mutated^4, 7-10^. Many ncMAPs are found exclusively in cancer cells and attract attention as (1) they can be immunogenic *in vitro* as well as *in vivo*; (2) they are more numerous in the immunopeptidome of malignant cells than mutated TAs and (3) several non-coding TAs are widely-shared between cancer patients whereas mutations mainly generate private antigens^11, 12^. In the context of proteogenomics usage, ncMAPs discovery and actionable TAs identification have raised three challenges that are often addressed inconsistently by immunologists.

First, the attribution of an exact RNA expression to MAPs. Typically, proteogenomic pipelines quantify MAPs RNA expression through the estimation of their parental transcript expression by using conventional transcript abundance quantification tools. However, such tools cannot be used reliably for ncMAPs which often derive from unannotated genomic regions. Furthermore, such approaches do not consider that MAPs (8-11 residues) could derive from multiple regions of the genome due to the degeneracy of the genetic code. Therefore, studies failing to consider all genomic regions susceptible to generating a given MAP would underestimate its RNA expression.

Second, the attribution of a biotype to MAPs. Due to the multiplicity of genomic regions able to generate the same MAP, and possibly having different biotypes, a MAP could be mislabeled for example as ERE-derived while a canonical region could also generate it through out-of-frame translation.

The third challenge is to prioritize TAs. Ideally, TAs should be immunogenic and specifically expressed (or overexpressed) by malignant cells^13^. Because RNA expression is a reliable proxy of the MAP presentation probability^8, 14^, RNA-seq data of tumor and normal samples are powerful tools to perform TA prioritization. While tumor specificity can be evaluated by comparing MAP RNA expression between tumor and normal samples, evaluating MAPs RNA expression in medullary thymic epithelial cells (mTECs) should be a good predictor of immunogenicity because mTEC MAPs induce central immune tolerance^15^. However, for the reasons mentioned above, comparing reliably MAPs RNA expression between tumors, their paired normal samples, and mTECs requires considering all their possible genomic regions of origin.

To address these challenges, we developed BamQuery, an annotation-independent tool that enables the attribution of an exhaustive RNA expression profile to any MAP of interest in any RNA-seq dataset of interest.

## 2. RESULTS

### Exhaustive capture of MAPs RNA expression

Because genomic annotations cover vast regions that are unlikely to represent accurately the local RNA expression of an 8-11 residues peptide (especially for ncMAPs deriving from introns, Extended Data Fig. 1a) and because no annotations are available for MAPs deriving from intergenic regions, we designed BamQuery to evaluate MAPs RNA expression independently of annotations. Due to the small size of MAP-coding sequences (MCS, 24-33 nucleotides), counting the RNA-seq reads containing each MCS able to code for a given peptide is the most thorough and less error-prone method to evaluate MAPs RNA expression. To make BamQuery readily available, it had to work on a broadly used data format. Given that querying MCS in fastq files is time-consuming (> 1 minute / MCS), we designed BamQuery to work on bam files in five steps (Fig. 1a, and Methods): (1) reverse-translation of each MAP into all possible MCS; (2) mapping of MCS to the genome using STAR^16^ to identify those having perfect matches with the reference and attribute them a genomic location. At this step, we also include to the reference genome the mutations from the dbSNP annotations^17^ to enable the mapping of mutated sequences; (3) counting of the primary RNA-seq reads encompassing exactly the MCS at their respective location (∼0.0005 minute / MCS / location) and sum read counts of each MAP across locations; (4) normalization of the read count of each MAP by the total primary alignment read count of the sample and multiplication by 1×10^8^ to yield read-per-hundred-million (RPHM) numbers and (5) attribution of biotypes to MAPs based on the reference annotations overlapping the various expressed (RPHM>0) regions.

**Fig. 1.**
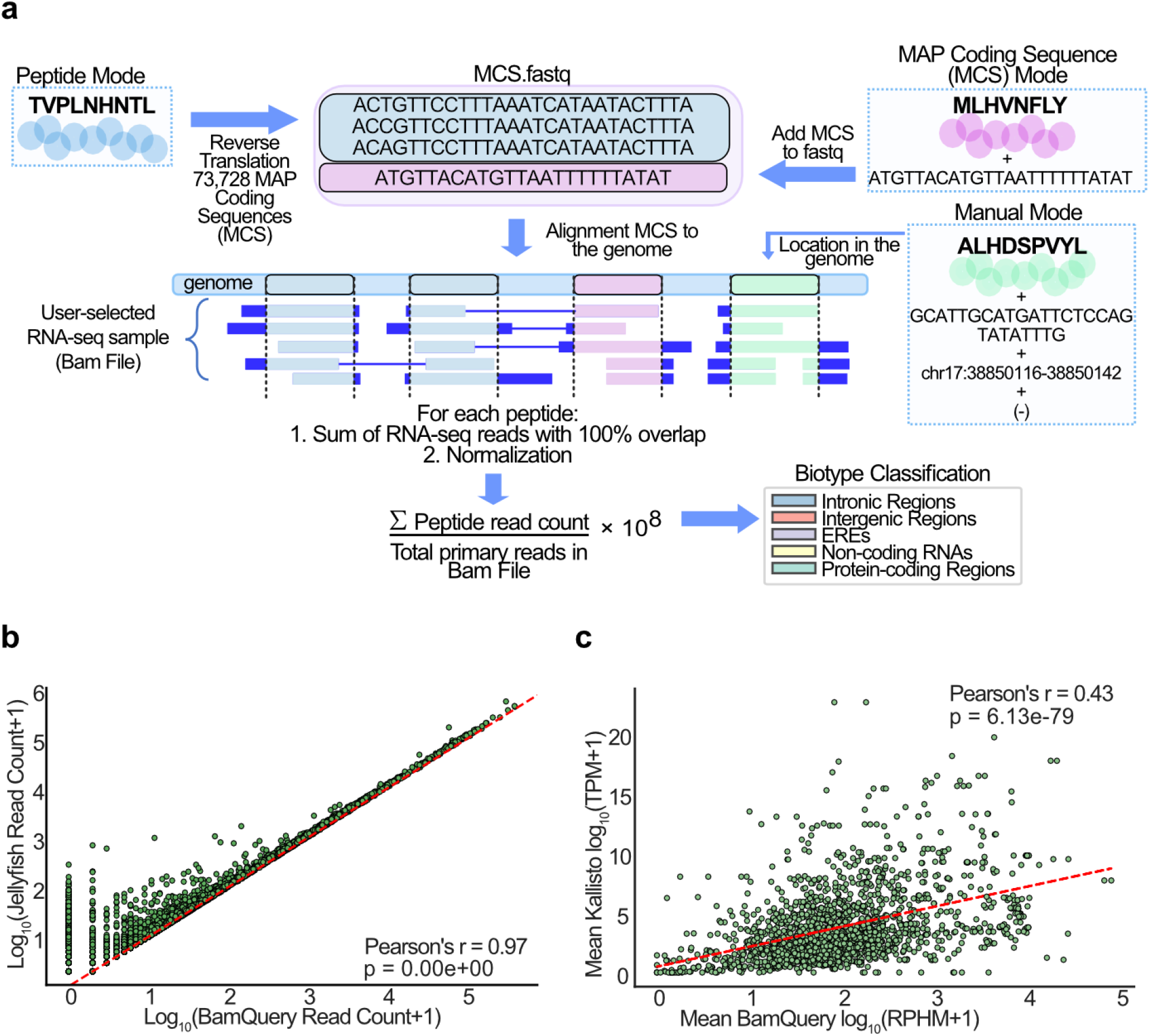
Exhaustive capture of MAPs RNA expression. **a**, Overview of the BamQuery approach to measuring MAPs RNA expression levels. **b**, Pearson’s correlation between BamQuery-acquired read counts and Jellyfish’s K-mer counts for MCS of canonical nine-mer MAPs (n=1,211) from the HLA Ligand Atlas (present in at least 20 different tissues) and 8 mTEC samples. **c**, Pearson’s correlation between BamQuery RPHM quantification and Kallisto TPM quantification for canonical MAPs (n=1,702) from the HLA Ligand Atlas (present in at least 20 different tissues) and 8 mTEC samples. Red lines in (b) and (c) are linear regressions.

To test BamQuery, we collected robustly validated benign MAPs from the HLA Ligand Atlas^18^ (1,702 canonical MAPs shared across at least 20 tissues, Extended Data Fig. 1b,c) and queried them in the transcriptome of eight mTEC samples sequenced previously^9, 19^. As a control, we used the primary reads contained in the mTEC bam files previously aligned with STAR to generate databases of 27-nucleotide-long k-mers (reads chunked into smaller sequences) and queried these databases for all possible 27-mer-MCSs encoding 9-amino acid-long MAPs (1,211/1,702). Despite rare discrepancies possibly due to limitations of the STAR aligner in BamQuery, the comparison of read counts between BamQuery and k-mer databases showed an almost perfect correlation (94% mean accuracy), demonstrating the exhaustivity of BamQuery (Fig. 1b, Extended Data Fig. 1d). Next, we compared BamQuery to Kallisto^20^, a transcript abundance quantification tool (reference MAPs RNA quantification method) that was chosen because it provides results similar to other tools while having the fastest computing speed^21^. A poor correlation between Kallisto and BamQuery was found (Fig. 1c) as most Kallisto measurements were skewed toward lower values than BamQuery’s. Specifically, Kallisto did not detect expression for 32 MAPs while BamQuery reported considerable RPHM values. In fact, BamQuery revealed that these MAPs are the result of multiple genomic locations (mean = 11) and are completely lost when only a single MAP source transcript is quantified (Extended Data Fig. 1e,f), as is typically done with transcript abundance quantification tools. Overall, these results evidence the accuracy and superiority of BamQuery over conventional approaches.

### New insights into the immunopeptidome biology

Next, we explored the biological features of the immunopeptidome by evaluating the expression of the 1,702 canonical MAPs from the HLA ligand atlas along with 724 MAPs previously reported as non-canonical (EREs, intronic, and ncRNAs-derived, Supplementary Table 1) in normal tissues, including mTEC samples^22^, and tissues from GTEx^23^ (Supplementary Table 2). BamQuery attributed a genomic location to 100% MAPs: among canonical MAPs, all originally annotated genes were attributed to their respective MAP by BamQuery and among a large list of well-annotated ncMAPs^8^, the originally annotated genomic location was re-located by BamQuery with an accuracy of 100%.

Comparing all 9-mers together (to prevent biases due to differences of length proportions), a higher number of possible MCS (total number of MCS after reverse-translation) was found for non-canonical vs canonical MAPs, especially for those mapping to introns and EREs (Fig. 2a). To better understand this bias, we investigated whether this could be linked to the degeneracy of codons. We found that residues encoded by six synonymous codons (R/L/S) were enriched in intron- and ERE-derived MAPs, with leucine being the most enriched (Fig. 2b-c). Previously, we observed that MAP source transcripts use rare codons more frequently than transcripts that do not generate MAPs^4^. Therefore, we hypothesized that ncMAPs would use rare codons more frequently than canonical ones. Indeed, we found that the genomic codon frequency of residues encoded by 6 synonymous codons (R/L/S) was on average lower than those encoded by lower numbers of synonymous codons (Extended Data Fig. 2a) and that the codons of ncMCS presented a lower genomic frequency than canonical ones (Fig. 2d). As rare codons are rate limiting for protein synthesis^24-26^ and as MAPs derive frequently from defective ribosomal products (DRiPs) generated by alterations of protein synthesis rate^27^, our data suggest that DRiPs contribute more to the generation of ncMAPs than to canonical ones.

**Fig. 2.**
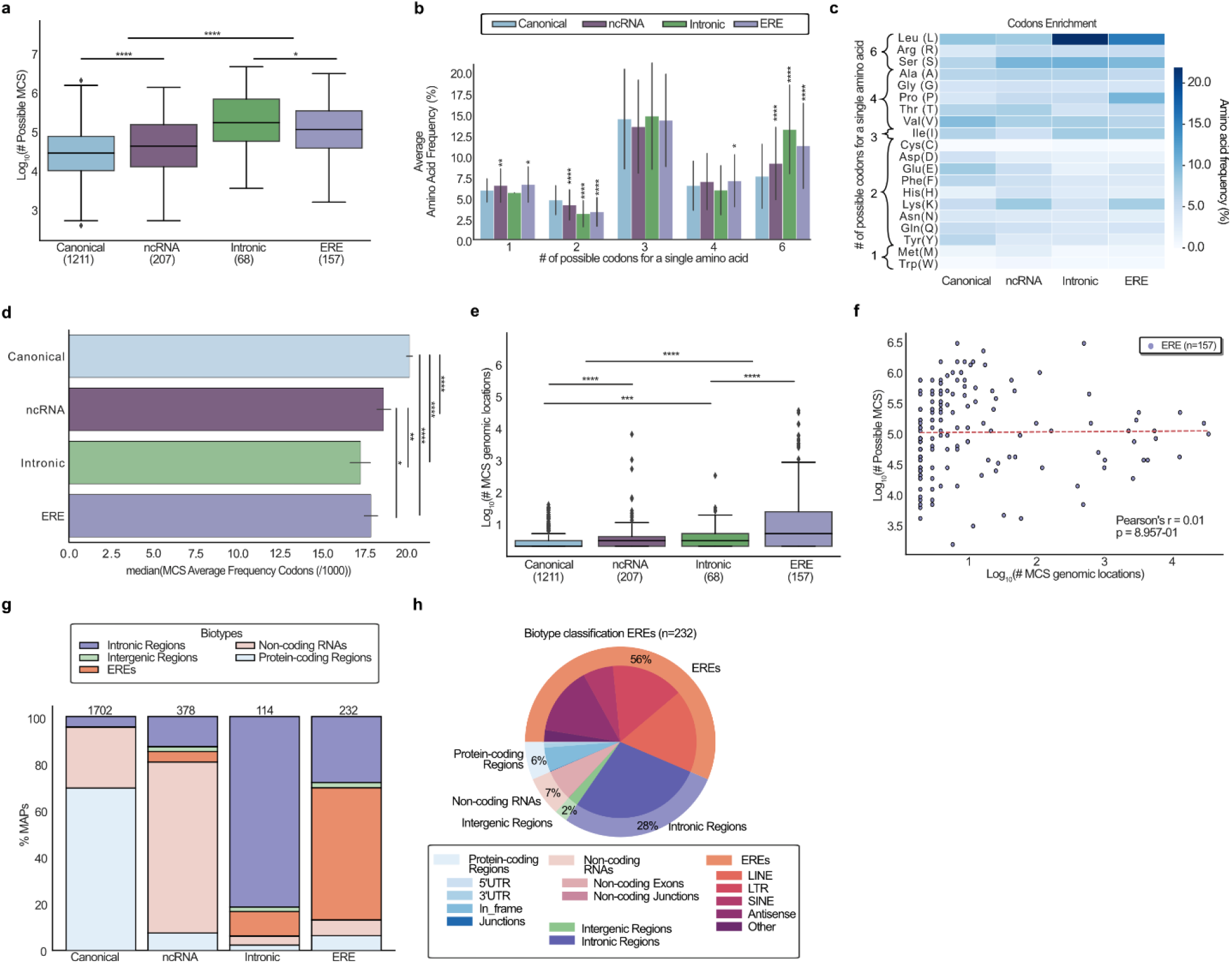
New insights into the immunopeptidome biology. **a-h** Published MAPs reported as canonical (n=1,702) and non-canonical (ncRNA (n=378), intronic (n=114) and EREs (n=232)) were searched with BamQuery in GTEx tissues and mTEC bam files in unstranded mode (GTEx data being unstranded) with genome version GRCh38.p13, gene set annotations release v38_104 and dbSNP release 151. Figures **a**,**e**,**f**,**g** were generated with the comparison of 9-mers only (n=1,211 canonical, n=207 ncRNA, n=68 intronic, n=157 EREs) to prevent possible biases introduced by variable frequencies of 8/10/11-mers among the compared groups. Figures **b**,**c**,**h** were generated with the complete MAP dataset (n=1,702 canonical, n=378 ncRNA, n=114 intronic, n=232 EREs). Mann-Whitney U test was used for indicated comparisons (*p<0.05, **p<0.01, ***p<0.001, ****p<0.0001). **a**, Number of possible MCS after reverse-translation of indicated MAP groups. **b**, Average frequency (%) of amino acids encoded by the indicated number of synonymous codons in indicated MAP groups. **c**, Heat map of amino acid frequency in indicated MAP groups. **d**, Mean of the MCS average usage frequency of codons (among 1000 codons located in human reference protein-coding sequences) encoding each of the 20 amino acids of indicated MAP groups. **e**, Number of MCS genomic locations able to code for the indicated MAP groups. **f**, Pearson’s correlation between the number of possible MCS after reverse translation vs the number of MCS genomic locations able to code for the assessed ERE MAPs. The red line is a linear regression. **g**, Percentage of MAPs attributed to indicated biotypes by BamQuery based on the EM-established biotype ranks and on the genomic regions expressed in GTEx tissues and mTECs. The X-axis indicates the biotype reported in the original study (groups). For clarity, BamQuery biotypes were summarized into five general categories: protein-coding regions, non-coding RNAs, EREs, intronic and intergenic. **h**, Percentage of the most likely biotype attributed by BamQuery to EREs MAPs.

Next, we analyzed the relation between the number of possible MCS per MAP (i.e., diversity of synonymous codons) and the number of genomic regions able to code for a given MAP. Canonical MAPs essentially derived from a single genomic location (60%), while non-canonical MAPs could derive from multiple regions (Fig. 2e). ERE MAPs presented the greatest numbers of possible regions, in agreement with their repeated nature (between 1.536 and 2.9×10^6^ possible regions). However, their number of possible MCS did not correlate with the number of possible locations, showing that amino acid residue composition cannot be used to predict the number of possible regions of origin (Fig. 2f).

Finally, given the multiplicity of possible regions of origin, we computed the most likely biotype of each MAP. For this, we used machine learning (expectation-maximization algorithm) to rank the biotypes (in-frame, intron, ERE, etc.) as a function of their likelihood of generating the reads covering them across the whole set of GTEx tissues. In general, canonical in-frame transcripts are more likely translated than non-canonical ones. For this reason, BamQuery’s best guess automatically ranks as “in-frame” any MAP having at least one in-frame canonical origin, which was the case for all canonical MAPs from our dataset (Extended Data Fig. 2b). BamQuery can also attribute biotypes based only on the likelihood ranks (considering the number of reads overlapping each transcript). In this case, ∼26% of canonical MAPs were assigned with a greater probability to ncRNAs (Fig. 2g). Furthermore, while ncRNA and intron MAPs were predicted to belong mainly from their identified biotype (73 and 81%) (Extended Data Fig. 2c), only 56% of ERE-derived MAPs were estimated to derive from EREs, and 6% of them could derive from canonical regions (5% in-frame) (Fig. 2h). Altogether, these data show that many published MAPs could be mislabeled, either as canonical or non-canonical.

### Single-cell proteogenomic analyses

High-throughput single-cell RNA sequencing (scRNA-seq) enables the examination of individual cells’ transcriptome^28, 29^. Therefore, we sought to perform single-cell analyses using BamQuery. Given the end-bias of the Chromium library design typically used in scRNA-seq, we evaluated whether read coverage would allow BamQuery analyses of canonical and non-canonical MAPs in cancerous^30^ and normal^31^ lung tissues scRNA-seq data. As expected, reads showed a bias toward the 3’ end of the canonical genes (Extended Data Fig. 3a). However, the coverage extended far from the 3’ end, in agreement with a report detecting mutations in various regions of the gene body^32^. We also found a surprisingly high (∼50% of reads) and homogeneous read coverage in introns and ERE regions, in agreement with previous reports^33, 34^, and suggesting that BamQuery would be able to detect expression for ncMAPs in scRNA-seq.

BamQuery detected expression for 50-60% of the canonical and non-canonical MAPs (Supplementary Table 1) in scRNA-seq, while 86% were found in bulk RNA-seq of GTEx lung samples (Fig. 3a). This lower number of MAPs expressed on single-cell data resulting from lower read coverage did not hamper the feasibility of scRNA-seq analyses. Indeed, canonical MAPs were uniformly expressed at the 5’ and 3’ ends of their transcripts (Fig. 3b). Also, the expressed rate of intronic and ERE MAPs in scRNA-seq data was more comparable to bulk RNA-seq data than canonical MAPs (Fig. 3c). This likely results from the more homogeneous read coverage observed in non-coding than in coding regions (Extended Data Fig. 3a).

**Fig. 3.**
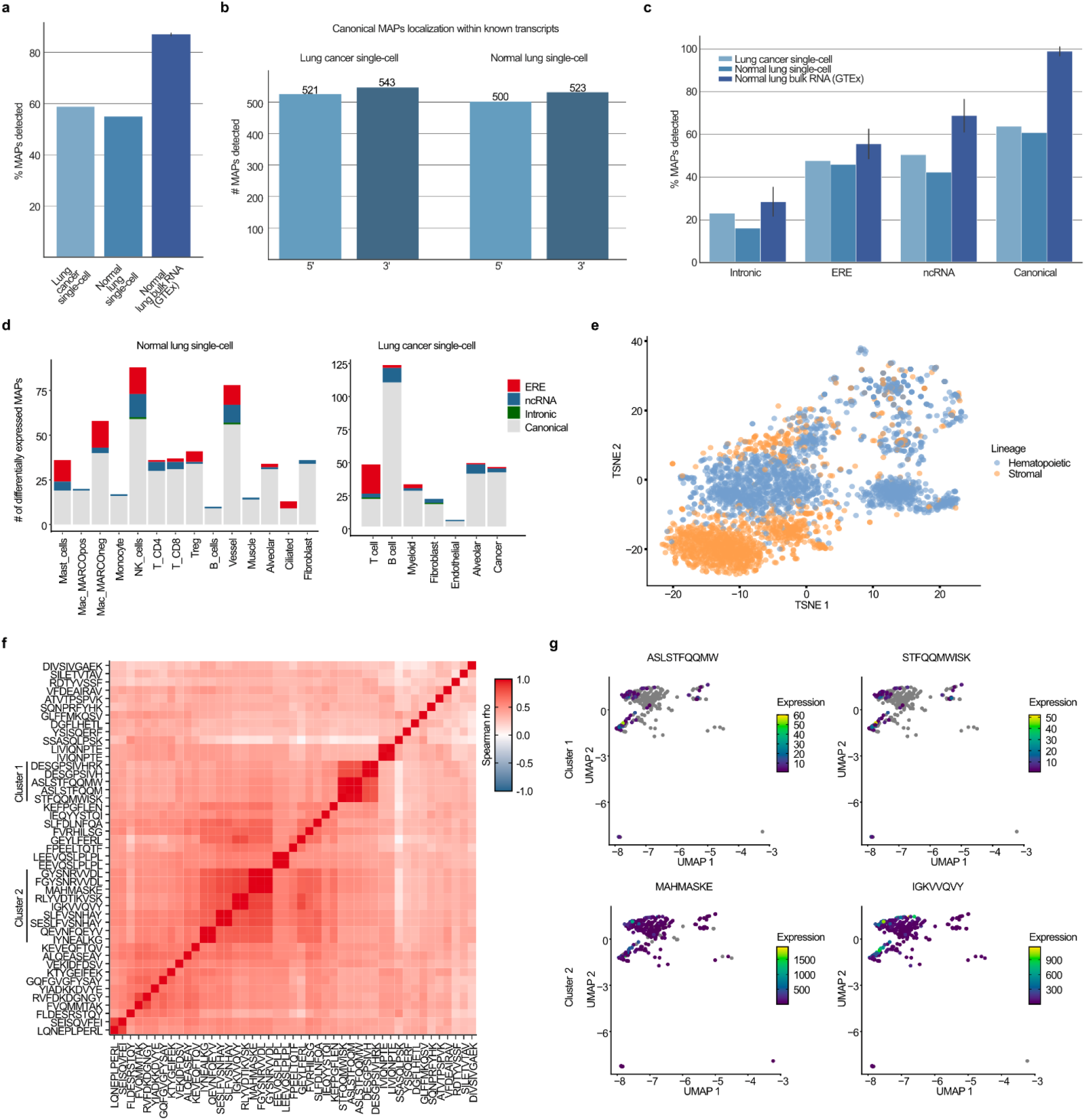
Single cell proteogenomic analyses. **a-g** Canonical (n=1,702) and non-canonical MAPs (ncRNA (378), intronic (114) and EREs (232)) were searched with BamQuery in bam files of scRNA-seq of normal and cancerous lung samples in single-cell in stranded mode with genome version GRCh38.p13, gene set annotations release v38_104 and dbSNP release 151. **a**, Median percentage of MAPs detected in normal and cancerous lung scRNA-seq, as well as in bulk RNA-seq samples of normal lungs from GTEx (n=150). **b**, Number of canonical MAPs located in the 5’ (first half of the transcript) or 3’ (second half of the transcript) region of the transcript detected in indicated scRNA-seq datasets. **c**, Median percentage of indicated MAP groups detected in normal and cancerous lung scRNA-seq, as well as in bulk RNA-seq samples of normal lungs from GTEx. **d**, Number of MAPs identified as differentially expressed by the different populations of cells in the normal lung (left panel) or cancerous lung (right panel). The originally reported biotype of the MAPs is indicated by the color code. **e**, TSNE analysis of the hematopoietic (blue) and stromal (orange) cells from the normal lung based on their MAP expression. **f**, Heatmap showing the co-expression (spearman rho, color bar) of MAPs overexpressed by lung cancer cells (rows vs columns). Two clusters of MAPs are highlighted on the left side of the heatmap (cluster 1 and cluster 2). **g**, TSNE showing the expression of MAPs (color bar) from cluster 1 (higher panel) or from cluster 2 (lower panel). Grey color indicates the null expression of a MAP in a cell.

Therefore, we explored the patterns of MAPs expression in normal and malignant lungs. Differential expression analysis showed that 12.86% (186/1446) and 16.46% (248/1506) of MAPs presented cell type-specific expression profiles in normal and malignant samples, respectively (Fig. 3d and Supplementary Tables 3-4). Several differentially expressed MAPs derived from genes having cell type-specific functions such as YTAVVPLVY in B cells (immunoglobulin J polypeptide), STFQQMWISK in muscle cells (Beta-actin-like protein 2), and FLLFPDMEA in macrophages (complement C1q B chain) (Extended Data Fig. 3b). To further assess the reliability of MAP expression, we re-clustered the normal lung dataset based uniquely on MAPs expression. This provided a clear separation of the hematopoietic and stromal compartments (Fig. 3e, Extended Data Fig. 3c) and allowed the clustering of specific cell populations such as alveolar cells or the monocytes and macrophages (Extended Data Fig. 3d,e). Strikingly, most MAPs identified as differentially expressed in the normal lung dataset had an expression restricted to either the hematopoietic or stromal lineages, showing a clear dichotomy between these two compartments in terms of MAP expression (Extended Data Fig. 3f).

Finally, given the growing interest in TAs shared between tumor cells, we assessed the clonality of 45 MAPs whose coding sequences were overexpressed by cancer cells through co-expression analyses. This highlighted two clusters of MAPs co-expressed in lung cancer cells (Fig. 3f) for which a distinct expression profile was observed in the lung (Fig. 3g). Indeed, MAPs of cluster 1 were expressed by a limited number of cancer cells, whereas MAPs of cluster 2 were ubiquitously expressed, making them more desirable immunotherapeutic targets. These data demonstrate the capacity of BamQuery to perform scRNA-seq analyses and evidence its potential to assess TAs intra-tumoral heterogeneity.

### MAP expression is underestimated in healthy tissues

Given the ability of BamQuery to capture MAPs RNA expression exhaustively, we evaluated the genomic origin of previously reported MAPs. First, we examined 1,062 colorectal cancer (CRC) TAs identified by their presence and absence from the immunopeptidome of malignant and paired benign cells, respectively^35^. To evaluate their probability of being presented by normal cells, we queried them in 3 datasets: GTEx, mTECs, and sorted dendritic cells (DCs)^36, 37^ (Supplementary Table 2). Four percent of TAs presented an expression <8.55 RPHM (minimum expression required to result in a probability >5% of generating a MAP^8^) in all normal tissues, except for testis, as these antigens would be classified as cancer-testis antigens (CTAs) (Fig. 4a). Strikingly, among the 7 TAs reported previously as being lowly expressed at RNA level in normal matched tissues, BamQuery revealed that only one (KYLEKYYNL) presented a low expression across all peripheral tissues. Finally, no expression was found for the RYLAVAAVF peptide (the only mutated TA reported in this study), while its wild-type counterpart was highly expressed, making it a promising target for CRC immunotherapies (Fig. 4b).

**Fig. 4.**
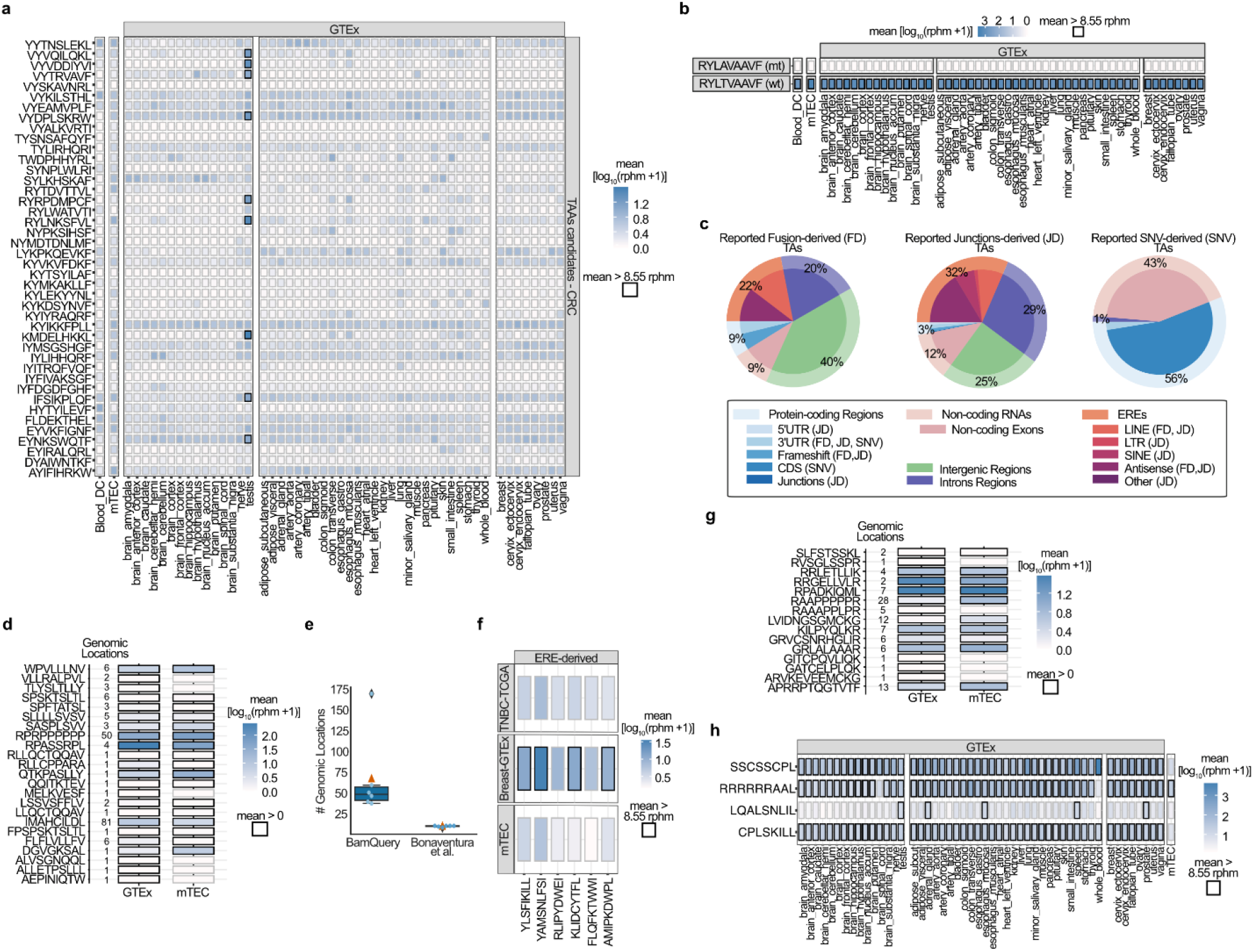
Underestimated MAP expression in healthy tissues. **a-g** Published human colorectal cancer (CRC) TAAs, mutated TAs, ERE-derived TSAs, proteasomal splicing peptides, and Epstein-Barr virus (EBV) MAPs were searched with BamQuery in the GTEx tissues (n=12–50 / tissue), mTECs (n=11) and/or DCs (n=19) bam files in unstranded mode with genome version GRCh38.p13, gene set annotations release v38_104 and dbSNP release 155 (except for the search for mutated TAs (**d**) where dbSNP was not considered, dbSNP=0). **a**, Heatmap of average RNA expression of published CRC TAAs in indicated tissues. Boxes in which a peptide has an average rphm>8.55 are highlighted in black. **b**, Heatmap of average RNA expression of the CRC mutated TA RYLAVAAVF and its wild type RYLTVAAVF in indicated tissues. **c**, Percentage of the most likely biotype attributed by BamQuery to published fusions, junctions, and SNVs-derived TAs. **d**, Heatmap of average RNA expression of published mutated TAs (n=23) in indicated tissues. The number of genomic locations expressed is presented on the left. **e**, Number of genomic locations at which the expression of the EREs TSAs was assessed by BamQuery vs by the original study. Light blue dots represent each assessed MAP and the orange triangle represents the average. **f**, Heatmap of average RNA expression of the EREs-derived TSAs in mTECs, normal breast tissues from GTEx (n=50), and triple-negative breast cancer samples from TCGA (n=158). **g**, Heatmap of average RNA expression of published proteasomal splicing MAPs (n=99) in indicated tissues. The number of genomic locations expressed is presented on the left. **h**, Heatmap of average RNA expression of EBV MAPs in indicated tissues.

Second, we wondered whether mutated TAs would be as tumor-specific as expected. We analyzed 45 8-11 amino acid long mutated peptides (7 from gene fusions, 28 from aberrant splice junctions, and 10 from single nucleotide variations, SNV) reported as tumor-specific in medulloblastoma (no RNA expression in GTEx)^38^. BamQuery could attribute a genomic location to 39 of them and mapped 7/10 SNV peptides to their reported genes (Extended Data Fig. 4a). Unexpectedly, BamQuery attributed non-discontinued (“unspliced”) expressed genomic locations to 82% of fusion and spliced peptides, evidencing that non-mutated (and mostly non-canonical, Fig. 4c) genomic regions could also code for those peptides. Overall, only 26 of 45 TAs presented low expression in normal tissues (Extended Data Fig. 4b) including all detected SNV-derived peptides. Therefore, we wondered whether mutated MAPs reported as cancer-specific in previous publications and public databases^10, 39, 40^ would be verified as such by BamQuery. From 323 mutated TAs (Supplementary Table 5), 23 (7%) were highly expressed in normal tissues where 25% of the peptides have more than 5 non-mutated genomic locations perfectly matching their MCS (Fig. 4d).

Third, we examined 6 ERE-derived MAPs reported as TAs (lowly expressed in normal tissues, including mTECs, and highly expressed in multiple cancer specimens) in triple-negative breast cancer^41^. While the original study identified an average of 8 locations for these peptides, BamQuery identified ∼66 locations per MAP (Fig. 4e). Moreover, these MAPs showed higher expression in normal breast samples compared to cancer samples (Fig. 4f). These results highlight the importance of considering all genomic locations able to generate a given MAP when measuring RNA expression.

Fourth, we evaluated whether BamQuery would detect non-discontinued genomic locations and RNA expression for MAPs supposedly impossible to be expressed by the human genome. We first examined 99 MAPs deriving from proteasomal splicing (generated from post-translational recombination of protein fragments)^42^. Fifteen could be generated by expressed regions (Fig. 4g), suggesting a possible misclassification of these peptides. Finally, considering the tight link between Epstein–Barr virus (EBV) infection and autoimmune disorders such as multiple sclerosis^43^, we examined the expression of 511 EBV-derived MAPs in the IEDB database. Four of them could be coded by the human genome and were expressed at high levels by normal tissues (Fig. 4h). Interestingly, one of them, CPLSKILL, can be presented by HLA-B8 molecules, an allele frequently associated with autoimmune disorders^44^.

Altogether, these results demonstrate that BamQuery is crucial to attribute an exhaustive RNA expression to MAPs and suggest that it could help select safe-to-target MAPs.

### Discovery of tumor-specific antigens in diffuse large B-cell lymphoma

Given the capacity of BamQuery to prioritize TAs, we wondered whether it could help identify tumor-specific antigens (TSAs) from raw immunopeptidomic data. By using a proteogenomic approach enabling the identification of TSAs^9^, we identified 6,869 MAPs from 3 published datasets of diffuse large B-cell lymphoma samples (DLBCL)^5^.

We first quantified the expression of the 6,869 MAPs in mTECs with BamQuery. A genomic location was found for 6,833 of them and most of them (∼86%) were highly expressed in mTECs (≥8.55 RPHM). To discriminate MAPs at risk of causing off-target toxicity when targeted, the remaining MAPs (14%) were queried in GTEx as well as in sorted benign B cells^36, 45^, and 5% of them were retained as being lowly expressed (<8.55 RPHM). Finally, the retained MAPs being upregulated (fold change ≥5) by the DLBCL samples in TCGA vs benign B cells and having evidence of translation based on the presence of ribosomal profiling elongation reads (queried with BamQuery in matched RIBO-seq data^5^, Extended Data Fig. 5a,b) were flagged as TSAs (67 MAPs, ∼1%, Fig. 5a, Supplementary Tables 6-7). Among them, 11 were promising as they were highly shared between DLBCL patients (Fig. 5b).

**Fig. 5.**
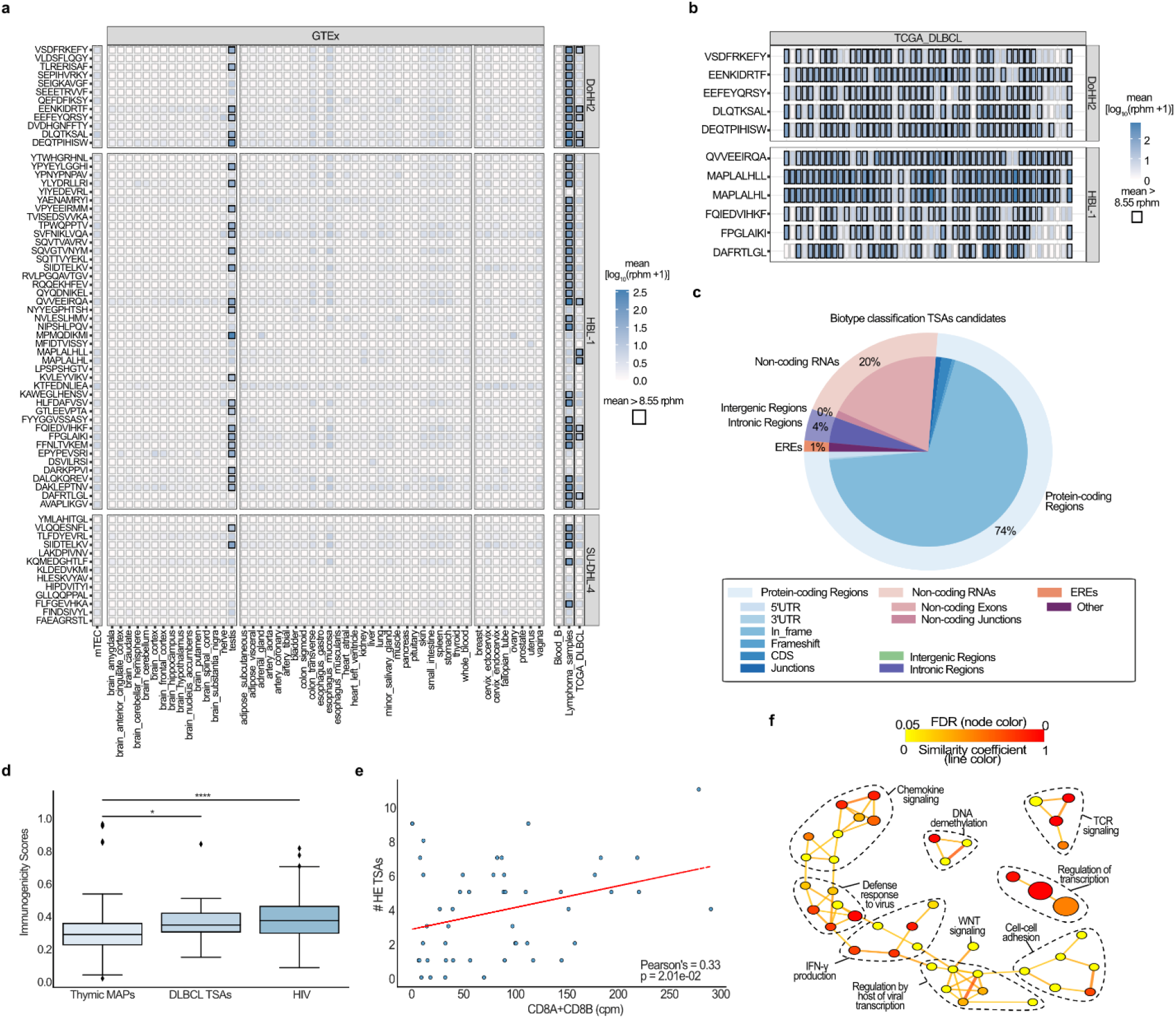
Discrimination of potential immunotherapeutic targets in DLBCL. **a-c** DLBCL MAPs, identified through a TSA-discovery proteogenomic approach, were searched with BamQuery in GTEx tissues (n=12–50 / tissue), mTECs (n=11), sorted blood B-cells (n=14), our DLBCL specimens (n=3) and/or TCGA DLBCL (n=48) bam files in unstranded mode with genome version GRCh38.104 and dbSNP version 155. **a**, Heatmap of average RNA expression of 67 TSA candidates in indicated tissues. Boxes in which a peptide has an rphm>8.55 are highlighted in black. **b**, Heatmap of average RNA expression of the highest shared and expressed TSA candidates (11) in cancer samples DLBCL from TCGA (n=48). Boxes in which MAPs expression (rphm) is >8.55 are highlighted in black. **c**, Percentage of the most likely biotype attributed by BamQuery for TSA candidates (n=67). **d**, Repitope immunogenic scores calculated for negative control thymic MAPs (n=158), highly expressed DLBCL TSAs (n=18, 25% of TSAs most upregulated by DLBCL TCGA versus normal blood in GTEx and sorted B cells), and positive control HIV MAPs (n=450). Mann-Whitney U test was used for comparisons (*p<0.05, ****p<0.0001). **e**, Pearson’s correlation in TCGA DLBCL patients (n=48) between the count of highly expressed (HE) TSAs expressed by each patient and the expression of cytotoxic T cells markers (CD8A+CD8B, in counts per million (cpm)). The red line is a linear regression. **f**, Network analysis of GO term enrichment among genes overexpressed by patients expressing an above-median number of HE-TSAs. Line color reflects the similarity coefficient between connected nodes. Node color reflects the false discovery rate (FDR) of the enrichment. Node size is proportional to gene set size.

BamQuery biotype classification showed that most TSAs derived from protein-coding regions of the genome, as only ∼25% of them derived from non-coding RNA (20%), EREs (1%), and intronic (4%) regions (Fig. 5c). Furthermore, based on their high expression in testis, 29 TSAs were flagged as CTAs^46^ (Supplementary Table 8) where most of them were known cancer biomarkers, supporting their relevance as immunotherapeutic targets. Additionally, upregulated TSAs in DLBCL samples compared to normal tissues (GTEx blood and benign B cells) had higher immunogenicity scores predicted by Repitope^47^ compared to previously published non-immunogenic controls^48^ (Fig. 5d). The expression of these TSAs correlated also with a greater expression of cytotoxic T cell markers (CD8A+CD8B), as well as with TCR signaling and other pro-inflammatory responses in DLBCL patients (Fig. 5e-f, Supplementary Table 9), supporting the biological value of TSAs discovered with BamQuery.

### BamQuery: an online tool to facilitate TA prioritization

We implemented an online portal to perform analyses on user-defined lists of MAPs. As we could not enable searches on GTEx (due to restricted use of these data), we included queries of MAPs in mTECs and DCs^36, 37^ (Supplementary Table 2) as a proxy of tumor-specificity and immunogenicity. While expression in mTECs is considered a good proxy for normal cell expression^49, 50^, we showed previously that mTECs share more transcriptomic features with epithelial than hematopoietic cells^8^. Prioritizing TAs based only on mTECs would therefore not be sufficient and we included DCs as they exert a non-redundant role in central tolerance establishment with mTECs^51^.

To validate this choice, we randomly selected 10% of hematopoietic-specific (2,429) and 10% of epithelium-specific (3,237) MAPs from the HLA ligand atlas (Extended Data Fig. 6a, b). We queried their expression in mTECs, DCs, GTEx epithelial tissues, and a set of hematopoietic cells (Supplementary Table 2). At the RNA level, DCs and mTECs presented the highest hematopoietic and epithelial MAPs expression levels, respectively (Extended Data Fig. 6c, d). We refined our analysis by focusing on hematopoietic and epithelial MAPs being lowly (<8.55 RPHM) expressed in mTECs and DCs, respectively. This revealed dramatically higher expression of hematopoietic and epithelial MAPs in hematopoietic (highest in DCs) and epithelial tissues (highest in mTECs), respectively (Fig. 6a, b). We conclude that MAPs lowly expressed in mTECs are highly expressed in DCs, and vice-versa.

**Fig. 6.**
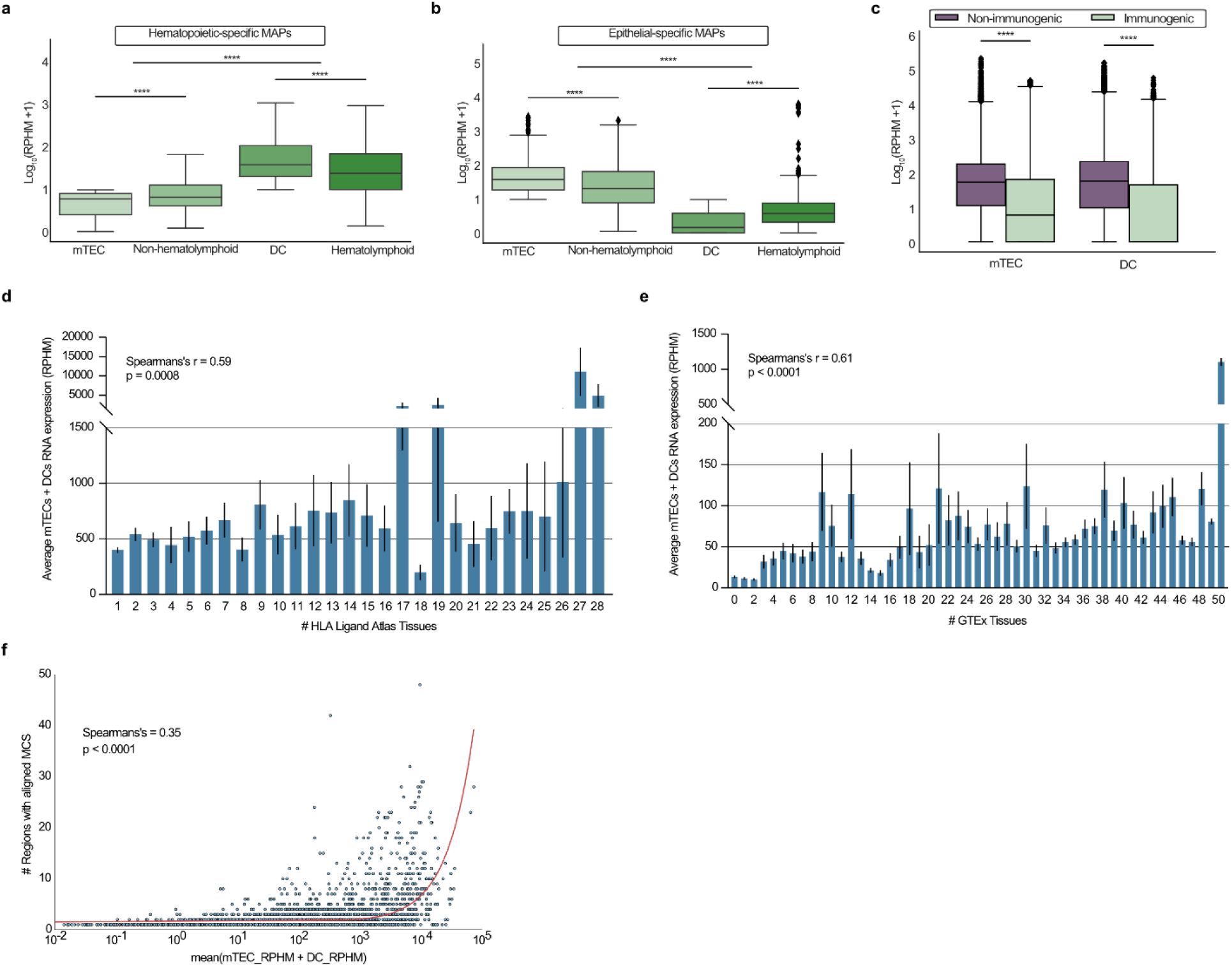
BamQuery: an online tool to facilitate TAs prioritization. **a-b**, Average RNA expression of hematopoietic-specific (**a**) and epithelial-specific (**b**) MAPs in mTECs (n = 11), non-hematolymphoid GTEx tissues (n = 2,389), DCs (n =19) and hematolymphoid GTEx tissues (n=196). Wilcoxon rank-sum test two-sided was used for comparisons (****p<0.0001). **c**, Average RNA expression of non-mutated human immunogenic (n=1180) and non-immunogenic (n=4917) MAPs in mTECs (n = 11) and DCs (n =19). Mann-Whitney U test was used for comparisons (****p<0.0001). **d-e**, Average mTECs+DCs RNA expression of a random selection of MAPs from the HLA Ligand Atlas (n=8621, 10% of the Atlas) as a function of the number of the HLA Ligand Atlas tissues presenting them (**d**) or as a function of the number of GTEx tissues in which the MAPs are expressed above an average of 8.55 RPHM (**e**). The average expression was correlated (Spearman) with the number of tissues. Error bar, SEM. **f**, Spearman’s correlation between the number of expressed genomic locations and the average expression in mTECs and DCs of the same MAPs used in (d). The red line is a linear regression (distorted by the log transformation of the x-axis).

Next, we tested whether MAPs expression in mTECs and DCs would predict their immunogenicity. We queried in mTECs and DCs 1,180 and 4,917 non-mutated human MAPs verified experimentally as immunogenic and non-immunogenic, respectively, and curated in Ogishi et al.^47^. Immunogenic MAPs presented a lower expression than non-immunogenic MAPs in both types of samples (Fig. 6c). On this dataset, we trained a logistic regression model to classify immunogenic and non-immunogenic MAPs using the RPHM values of mTECs and DCs as features. Measurements of model performance and robustness using the cross-validation method (area under the ROC curve (AUC) = ∼0.75, Extended Data Fig. 6e) showed that the RPHM values of MAPs in mTECs and DCs are predictors of MAP immunogenicity.

Finally, we evaluated whether MAP expression in both mTECs and DCs reflects the probability of presentation in benign tissues. Using 10% of random MAPs from the HLA Ligand Atlas (8,694), we found that MAPs lowly expressed in both mTECs and DCs were less presented (Fig. 6d) and expressed (Fig. 6e) in tissues of the HLA Ligand Atlas and GTEx, respectively. Upon examination of these MAPs features, we found that the probability of being highly expressed in mTECs and DCs increased exponentially with the number of possible genomic regions (Fig. 6f). Altogether, these results show that concomitant expression in mTECs and DCs expression is a reliable proxy of the presentation/expression in benign tissues and that MAPs having fewer possible regions of origin have a greater probability of being safe-to-target TAs.

The BamQuery public interface is accessible through http://bamquery.iric.ca/ and incorporates the logistic regression predictor model to report the conferred probability that a MAP is immunogenic. BamQuery is also available as a standalone version that can be configured to work with proprietary bam files. We believe that BamQuery will greatly help researchers in their attempts to identify specific and immunogenic TAs.

## 3. DISCUSSION

Fuelled by studies focused on TAs, the immunopeptidomics field is expanding rapidly^3, 52, 53^. This expansion comes with an impressive diversity of homemade methodological approaches addressing the challenges raised by the characterization of non-canonical and mutated MAPs. Specifically, the fact that ∼75% of the human genome can be transcribed^54^ (and therefore possibly translated) evidenced the necessity of examining the expression of each region able to code for a presumed TA. BamQuery was designed not only to enable such examination but also to enable a uniformization of TA validation approaches across laboratories.

The recent discovery that a significant fraction of the immunopeptidome derives from non-coding regions has brought the contribution of the “dark genome” into the spotlight^2^. Since then, multiple studies have attempted to characterize cryptic MAPs, most often by using mass spectrometry informed by databases dedicated to the identification of specific classes of ncMAPs (intron-derived, ERE-derived, etc.)^7, 19, 55^. However, these approaches suffer from their dedication as the identified MAPs could also derive from other transcripts, absent from these databases. Accordingly, based on evidence showing that greater RNA expression confers a greater probability of MAPs generation^7, 13^, we implemented a biotype annotation tool in BamQuery and showed that many presumed ncMAPs could be coded with greater probability by regions annotated with different biotypes. Strikingly, an important fraction of the canonical MAPs (∼30%) could also be translated, with a greater probability, from non-canonical regions. While this result requires more in-depth analyses to elucidate the true origin of these MAPs, this possible dramatic contribution of the non-coding genome to the immunopeptidome is a sobering thought given that cryptic proteins are translated as efficiently as canonical proteins and generate MAPs 5-fold more efficiently per translation event^5^.

Therapies targeting truly tumor-specific antigens can be highly effective^56^, while those targeting antigens unsuspectedly expressed by normal cells can be lethal for patients^57^. Notably, BamQuery evidenced a high expression of many TAs, including mutated and ERE MAPs, in normal tissues, resulting from previously unreported coding regions and suggesting that targeting them would be unsafe. Here, we acknowledge that our approach can be considered very cautious. Indeed, by summing the RNA-seq reads of all regions able to code a given TA, BamQuery does not consider that possibly only one region is translated and generates MAPs. Eventually, the availability of RIBO-seq data (which can be analyzed with BamQuery as well) could help to address this question. Meanwhile, in the absence of tools robustly predicting the translational origin of MAPs, the approach reported herein is the most circumspect for TA selection. Ideally, we recommend prioritizing TAs with a single possible region of origin (with cancer-specific expression) because other regions cannot code for such TAs in normal tissues.

Thanks to its exhaustivity, speed, ease of use, and versatility (bulk & single-cell RNA-seq + RIBO-seq, usable with mouse or human genome, on any kind of wild-type or mutated MAPs), BamQuery enables for the first time a uniformization of proteogenomic analyses in MHC-I immunopeptidomics.

## Acknowledgments

We are grateful to Qingchuan Zhao, Assya Trofimov, Nandita Noronha, and Caroline Labelle for useful biological insights, suggestions, and testing BamQuery. We also thank all other members of our laboratories for their thoughtful recommendations. We thank Eric Audemard and Geneviève Boucher of the IRIC bioinformatic platform for assistance with bioinformatics tools. This study was supported by grants from the Canadian Cancer Society (707264), and the Canadian Institutes of Health Research (FDN 148400). GE is supported by post-doctoral fellowships from the IRIC, FRQS, The Cole Foundation, and the FNRS. We thank the Genotype-Tissue Expression (GTEx) Project for providing RNA-seq data. The GTEx Project was supported by the Common Fund of the Office of the Director of the National Institutes of Health, and by NCI, NHGRI, NHLBI, NIDA, NIMH, and NINDS.

## Author Contributions

MVRC coded the final version of BamQuery, designed the study, performed the analyses, interpreted the data, and wrote the article. JDL performed the single-cell analyses. MPH, AA, EK, and KV helped with bioinformatic analyses and data interpretation. PG deployed the web portal and prepared BamQuery docker. JPL generated DLBCL cell lines MS databases. CD performed MS searches for DLBCL samples. GE coded the first version of BamQuery and helped with data analyses. GE, CP, SL, and PT designed the study, interpreted the data, and wrote the manuscript. All authors edited and approved the manuscript.

## 4. METHODS

### Data and Code Availability

The Python and R scripts generated during this study are available on GitHub, https://github.com/lemieux-lab/BamQuery. The standalone version of BamQuery can be downloaded at http://bamquery.iric.ca/installation.html. Details regarding all samples used in this study are listed in Supplementary Table 2.

### Datasets

The eight human mTEC samples have been prepared and sequenced for previous studies of our team (GEO:GSE127825 and GEO:GSE127826) (Larouche et al., 2020^19^; Laumont et al., 2018^9^). Three additional mTEC samples were published (ArrayExpress:E-MTAB-7383) by Fergurson et al. ^22^.

### BamQuery

BamQuery is designed to analyze MAPs ranging in length from 8 to 11 amino acids (aa). As peptide input, BamQuery supports three different formats that can be pulled into a single input file.

A) Peptide mode: only the amino acid sequence of the MAP is provided, hence BamQuery performs a comprehensive search for its RNA-seq expression. All results reported in the present article were obtained with this mode.

B) MAP coding sequence (MCS) mode: the amino acid sequence of the MAP is provided, hence BamQuery performs the search for the expression of the given MCS only.

C) Manual mode: the amino acid sequence of the MAP is provided followed by an MCS, the corresponding location in the genome of the given MCS, and the strand (+ forward or - reverse), whereby BamQuery performs the expression search at the given location for the given MCS at the given genomic location and strand (useful to evaluate the expression of mutated MAPs whose genomic location is known but which cannot be located by BamQuery due to unavailable annotations in dbSNP or STAR failure).

BamQuery performs five important steps for each peptide queried.

#### 1) Reverse translation of MAPs

Each input MAP in peptide mode is reverse-translated into all possible MCS. The MCS are compiled into a fastq file.

#### 2) Identification of genomic locations

MCS are then mapped to the reference genome (user-defined, meaning that several genome versions are supported (GENCODE 26, 33 or 38)) using STAR v2.7.9.a^16^ running with default parameters except for –seedSearchStartLmax, --winAnchorMultimapNmax, -- outFilterMultimapNmax, --limitOutSJcollapsed, --limitOutSAMoneReadBytes, -- alignTranscriptsPerWindowNmax, --seedNoneLociPerWindow, --seedPerWindowNmax, -- alignTranscriptsPerReadNmax that were replaced by 20, 10.000, 10.000, 5.000.000, 2.660.000, 1.000, 1.000, 1.000, 20.000, respectively. MCS genomic locations (perfect alignments) are selected from the output STAR file Aligned.out.sam. Perfect alignments are defined as MCS matching exactly the reference genomic sequence or as MCS bearing mismatches annotated as known polymorphisms in the dbSNP database (user-selected dbSNP 149, 151, or 155 releases). Therefore, each alignment included in Aligned.out.sam is exanimated to compare the read sequence nucleotide by nucleotide against the reference genomic sequence at that position (assessed using the samtools fetch command within python via the pysam (https://github.com/pysam-developers/pysam) library at the genomic location of the given alignment). If a difference is detected between a nucleotide of the aligned read sequence and the nucleotide of the reference genomic sequence at a given position, the position is queried in the python dictionary containing the SNVs of the dbSNP database selected by the user. If all discrepancies in the current alignment are known (supported by the SNVs in the dbSNP database) the alignment is retained as it is considered perfect, otherwise, the alignment is discarded. To reduce the complexity of tracing perfect STAR alignments, only single nucleotide variants (SNVs) of dbSNP annotations were considered to define perfect alignments.

#### 3) MAP RNA-seq reads counting

Next, the expression of each MCS is queried in each BAM file (CRAM files are also supported) using the samtools view^58^ command within python via the pysam library (only primary alignment reads (pysam option -F0X100), originally present in fastq files, are queried) at their respective genomic location. BamQuery supports RNA-seq unstrandedness / strandedness libraries (user-defined parameter, default: strandedness). To collect reads in unstranded libraries, the -F0X100 option is used in the pysam view command. In stranded libraries, depending on the sequencing reads type (single-end, paired-end), library preparation (forward or backward) and sense of the MCS genomic location (forward or backward), the options in the pysam view command are: - F0X100 & -f0X50 for R1 mate and -F0X100 & -f0XA0 for R2 mate in paired-end, forward library and reverse genomic location; -F0X100 & -f0X60 for R1 mate and -F0X100 & -f0X90 for R2 mate in paired-end, forward library and forward genomic location; -F0X110 for R1 mate in single-end, forward library and forward genomic location; -F0X100 & -f10 for R1 mate in single-end, forward library and reverse genomic location; -F0X100 & -f0X60 for R1 mate and -F0X100 & -f0X90 for R2 mate in paired-end, reverse library and reverse genomic location; -F0X100 & - f0X50 for R1 mate and -F0X100 & -f0XA0 for R2 mate in paired-end, reverse library and forward genomic location; -F0X110 for R1 mate in single-end, reverse library and reverse genomic location; -F0X100 & -f10 for R1 mate in single-end, reverse library and forward genomic location. The retrieved reads are examined one by one and counted if they exactly span the queried MCS at the genomic location. Therefore, each retrieved read is transformed into a list in Python and its alignment location is transformed into an array containing the location of each amino acid in the read. The indices of the array locations corresponding to the first and last amino acid locations in the MCS at a given genomic location are used to extract from the read list the subsequence that is compared to the MCS. If both the MCS and the subsequence of a retrieved read are the same, the read count for the current MCS increases by one. Finally, the total read count (tr_MAP_) for a given MAP is computed by summing all RNA-seq reads from all MCS genomic locations.

#### 4) Normalization

The tr_MAP_ count is transformed into “reads per hundred million” values (RPHM) by normalizing them with the total number of primary reads sequenced (corresponding to the total read number present in fastq files) according to the formula: 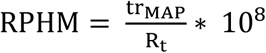 where R_t_ represents the total number of primary RNA-seq reads of the sample. These final values are log-transformed log_10_(RPHM + 1) to allow comparison and averaging between samples, thus removing the bias of large values.

#### 5) Biotype classification

All genomic locations identified for each MAP are compiled into a bed file and their biotypes are obtained using BEDtools^59^ intersect with the following options -a (annotation file), -b (genomic locations), -wao (writes the original annotation and genomic location entries along with the number of base pairs of overlap between the two features), and the following annotations: RepeatMasker (GRCh38/hg38 assembly, to annotate the EREs) and GENCODE (for all other biotypes, gene set annotations releases v26_88, v33_99, v38_104). The complete list of biotypes annotated by BamQuery based on RepeatMasker and GENCODE can be consulted at http://bamquery.iric.ca/biotype_classification.html.

Given that MAPs may have alignments in regions where several different biotypes overlap (such as protein-coding transcripts overlapping with non-coding RNAs, see the example shown in Extended Data Fig. 7a), we used the expectation-maximization (EM) statistical model to estimate, for each biotype, the read distribution coefficient. In this model, reads at each genomic location are weighted for each biotype at the given location according to their coefficients and consequently, the biotype of each MAP is scored according to the percentage of reads corresponding to each biotype (in-frame, introns, ncRNA, ERE, etc.). The EM algorithm iterates between the expectation (E) and maximization (M) step until the parameter set of the last iteration is unchanged, therefore finding the parameter set that maximizes the posterior probability of the observed data, in our case the reads that overlap with one or more biotypes. To train the EM algorithm, we first collected canonical and ncMAPs (Supplementary Table 1) and ran BamQuery on normal and cancer datasets (normal: GTEx and mTECs, cancer: TCGA) to obtain the total reads covering each MAP at each MCS genome location. We then computed the probability of each biotype as follows:

Let ∅ = (∅_A,_ ∅_B_, ∅_c_ …), be the set of parameters to estimate, where ∅_A,_ ∅_B_, ∅_c_… are the probabilities that the read belongs to the In_frame (A), non_coding_exon (B), intron (C), etc. biotypes. EM starts with an arbitrary initial estimation of 0.1 for each biotype’s probability. In the E-step, the distribution of the total number of reads for each MAP is computed using the current biotype’s parameters, as follows:

Let R_i_ = total reads of MAP_i_

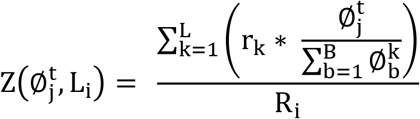

Where 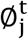 is the current probability for biotype j in MAP_i_. L_i_ is the MCS genome locations for MAP_i_. r_k_ is the number of reads overlapping location k and B is the set of biotypes overlapping the location k.

In the M-step, the new set of parameters is determined using the current computations, as follows:

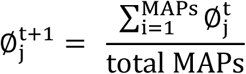

Where 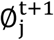 is the new probability for biotype j obtained after summing all the probabilities distributions of all MAPs computed in the last E step and normalizing by the total number of MAPs. The iterative process concludes if the following condition is met for all biotypes: 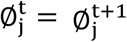 and the last set of estimated parameters is used to assign the proportion of reads assigned to each biotype at any genomic location.

Therefore, BamQuery scores for each MAP the biotype as the percentage of reads assigned to each biotype class (in-frame, introns, ncRNA, ERE, etc.). For example, a canonical MAP with alignments in non-canonical regions could be indicated as follows In_frame: 84.09% - Intronic: 15.91%, meaning that ∼84% of the total reads overlap with a known transcript and that the MAP is within the known protein frame, while ∼16% of the reads overlap with transcripts in an intronic region.

BamQuery informs the biotype of each MAP in three different settings, as follows:

1. Biotype computed for each MCS genome location: BamQuery reports the percentage contribution of the biotypes overlapping the given location. The percentage of each biotype is calculated as the coefficient of each biotype normalized by the sum of the coefficients of all biotypes in the location, as follows:

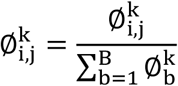

Where 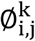 is the coefficient assigned to the biotype j for the MAP_i_ at the location k.
2. Biotype computed from all MCS genome locations found in the set of queried samples: the biotype of each MAP is assigned based on the total read count in the sample set. This calculation follows three steps:
  a. The total number of reads in each MCS genome location is distributed according to the biotype percentages assigned to the location in the previous step.
  b. Normalization of the distributed count of reads by the total number of reads in the entire set of samples.
  c. The final biotype of each MAP is obtained by summing all normalized reads distributions across its MCS genomic location.
3. Biotype for each subset of samples (e.g., GTEx, TCGA, mTEC samples): the biotype of each peptide is assigned following the same steps as before but according to the total count of reads in each subset of samples.
4. Best guess biotype: BamQuery also reports the most likely biotype for each MAP (Best Guess) following the rules below:
  a. Since a MAP is most likely to be generated from a known canonical protein if the MAP ever appears in-frame of a protein the best guess assigned is In-frame with the certainty given in the biotype classification.
  b. Otherwise, the best guess biotype is assigned according to the biotype with the highest percentage of the biotype ranking.

Full documentation of supported options, examples of use, and descriptions of BamQuery reports can be found at http://bamquery.iric.ca/

### K-mer databases

K-mer databases were generated by retrieving the primary mapped reads from the bam files of each mTEC sample with samtools view ^58^ (-F260 option) followed by SamToFastq from Picard tools to recover R1 and R2 fastq files (https://broadinstitute.github.io/picard/index.html). Next, R1 reads were reverse complemented using the fastx_reverse_complement function of the FASTX-Toolkit v0.0.14. and fastq files of all mTEC samples were concatenated. Finally, Jellyfish count (v2.2.3, options -m = 27 and -s =1G)^60^ was used to generate the database from the fastq file, and jellyfish query was used to query the MCS in the database.

### Kallisto quantification

Transcript expression quantifications of mTEC samples were performed with kallisto^20^ v0.43.0 quant with default parameters except for --rf-stranded. The expression of each HLA atlas peptide was obtained from the mean TPM expression value of all transcripts associated with the peptide source genes.

### BamQuery Accuracy

For each MCS of the canonical nine-mer MAPs, we defined the BamQuery accuracy, as follows:

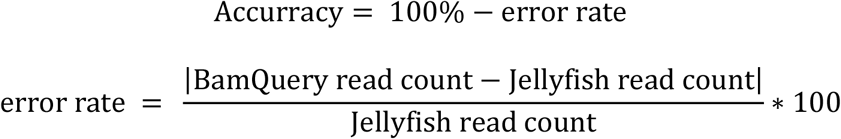

Therefore, the accuracy is the difference in the error rate with respect to 100%, the error rate being the percentage value of the difference in the observed MCS read count in BamQuery and the actual MCS read count in Jellyfish.

### Single cell RNA-seq analyses

Previously published single-cell RNA-seq data from the healthy and cancerous lungs were downloaded from the NCBI BIOPROJECT (accession number PRJEB31843) and Array Express (accession number E-MTAB-6653), respectively. Reads were aligned on the human reference genome (GRCh38) using STAR version 2.7.9a^16^. Cell population annotations were performed using gene lists from Madissoon et al.^31^ and Lambrechts et al.^30^ for the healthy and cancerous lung datasets, respectively. For the subsequent profiling of MAP expression with BamQuery, the HCATisStab7509734 and the BT1375 samples were subsampled from the healthy and cancerous lung datasets, respectively. Read count normalization, log transformation, and dimensionality reduction were performed using the scater v1.18.6 and scran v1.18.7 R packages. The differential expression analyses of MAPs between the cell populations of the healthy and cancerous lungs were performed with the FindAllMarkers function of Seurat with the MAST model. Cells of the healthy lungs were also re-clustered based on their MAP expression using the runUMAP and runTSNE functions of the scater package, and cell lineages and populations previously annotated based on gene expression were represented on the resulting UMAP and TSNE graphs. Co-expression of MAPs in the tumor cells of the lung was also assessed. To do so, we selected the MAPs identified as overexpressed in lung cancer cells by the differential expression analysis and computed spearman correlations between the expression of each possible pair of MAPs. Finally, MAP expression in the cell populations of the healthy lung was visualized with Seurat^61^ v4.1.0.

### Immunogenicity predictions

Immunogenicity predictions of HE-TSAs were performed with Repitope^47^. Feature computation was performed with the predefined MHCI_Human_MinimumFeatureSet variable and updated (July 12, 2019) FeatureDF_MHCI and FragmentLibrary files provided on the Mendeley repository of the package (https://data.mendeley.com/datasets/sydw5xnxpt/1). HIV MAPs (positive control) were obtained from https://www.hiv.lanl.gov/content/immunology/tables/ctl_summary.html.

### Differential gene expression analysis

Transcript expression quantifications were performed on TCGA DLBCL bulk RNA-seq samples with kallisto v0.43.0 with default parameters. Then, with BamQuery, we attributed to each patient a count of highly expressed TSA transcripts (HE-TSA), i.e., the number of TSAs whose expression was above their median RNA expression across all patients having a non-null expression of the given TSAs. Patients having an above-median number of HE-TSAs (n=26) were compared to those below-median (n=22) through a differential gene expression analysis. This analysis was conducted in R3.6.1 as reported previously^62^. In brief, raw read counts were converted to counts per million (cpm), normalized relative to the library size, and lowly expressed genes were filtered out by keeping genes with cpm >1 in at least 2 samples using edgeR 3.26.8 and limma 3.40.6. This was followed by voom transformations and linear modeling using limma’s lmfit. Finally, moderated t-statistics were computed with eBayes. Genes with p-values < 0.05 and -1≥log_2_(FC)≥1 were considered significantly differentially expressed (386 genes upregulated and 1304 downregulated).

### GO term and enrichment map analyses

Biological-process gene-ontology (GO) term over-representation was performed with DAVID (https://david.ncifcrf.gov) on genes upregulated by DLBCL patients expressing high levels of HE-TSAs. Functional annotations with p-value < 0.05 were considered significant. The GO-term list was then imported in Cytoscape v3.7.2 and used to cluster redundant GO terms and visualize the results with EnrichmentMap v3.2.1 and default parameters. The network was visualized using the default “Prefuse Force-Directed Layout” in Cytoscape. Groups of similar GO terms were manually circled.

### Other bioinformatic analyses

Amino acid compositions were assessed with the ProtParam module of Biopython. Read coverage in scRNA-seq data was evaluated with the geneBody_coverage module of RSeQC on the bam file generated by CellRanger. Codon frequencies were obtained from the codon usage database (http://www.kazusa.or.jp/codon/).

### Logistic regression model

The cross-validation procedure was used to split the training data set into training and validation subsets using the StratifiedShuffleSplit function of the sklearn python library with 10 numbers of splits and 0.2 for test size. Next, the logistic regression model of the sklearn python library was used to classify immunogenic and non-immunogenic MAPs with the default parameters except for the liblinear solver.

### Construction of MS database for TSA identification

We used RNA-seq data from 3 published datasets of diffuse large B-cell lymphoma samples (DLBCL)^5^. Cancer-specific proteomes were built using k-mer profiling as described previously^9^. RNA-Seq reads were chopped into 33-nucleotide k-mers and only those present <2 in mTECs were kept. Overlapping k-mers were assembled into contigs, which were then three-frame translated and linked using “JJ” as separators. This database was concatenated with each sample’s canonical proteome for MAP identification.

### Quantification and Statistical Analysis

All statistical tests used are mentioned in the respective figure legends. For all statistical tests, *, **, ***, *** and **** refers to p< 0.05, p< 0.01, p< 0.001 and p< 0.0001, respectively, and are reported in the figures. Correlations were assessed with the Pearson or Spearman correlation coefficient, a red line in the correlation plots represents the linear regression. Plots and statistical tests were performed using scipy.stats and seaborn packages of Python v3.6.8. Unless mentioned otherwise, all boxes in box plots show the third (75th) and first quartiles (25th) and the box band shows the median (second quartile) of the distribution; whiskers extend to 1.5 times the interquartile distance from the box. Unless mentioned otherwise, all bar plots show the average with error bars: 95% confidence interval (CI).

## ABBREVIATIONS

MAP: MHC-I associated peptide
TA: Tumor antigen
ncMAP: non-canonical MAP
ncRNA: non-coding RNA
ERE: Endogenous retroelement
MCS: MAP coding sequence
ncMCS: non-canonical MCS
RPHM: Read-per-hundred-million
mTEC: Medullary thymic epithelial cell
DC: Dendritic cell
TSA: Tumor-specific antigen
CTA: Cancer-testis antigen
DLBCL: Diffuse large B-cell lymphoma

