## Supplementary Figures for "BamQuery: a proteogenomic tool for the genome-wide exploration of the immunopeptidome"

### Supplemental Figures

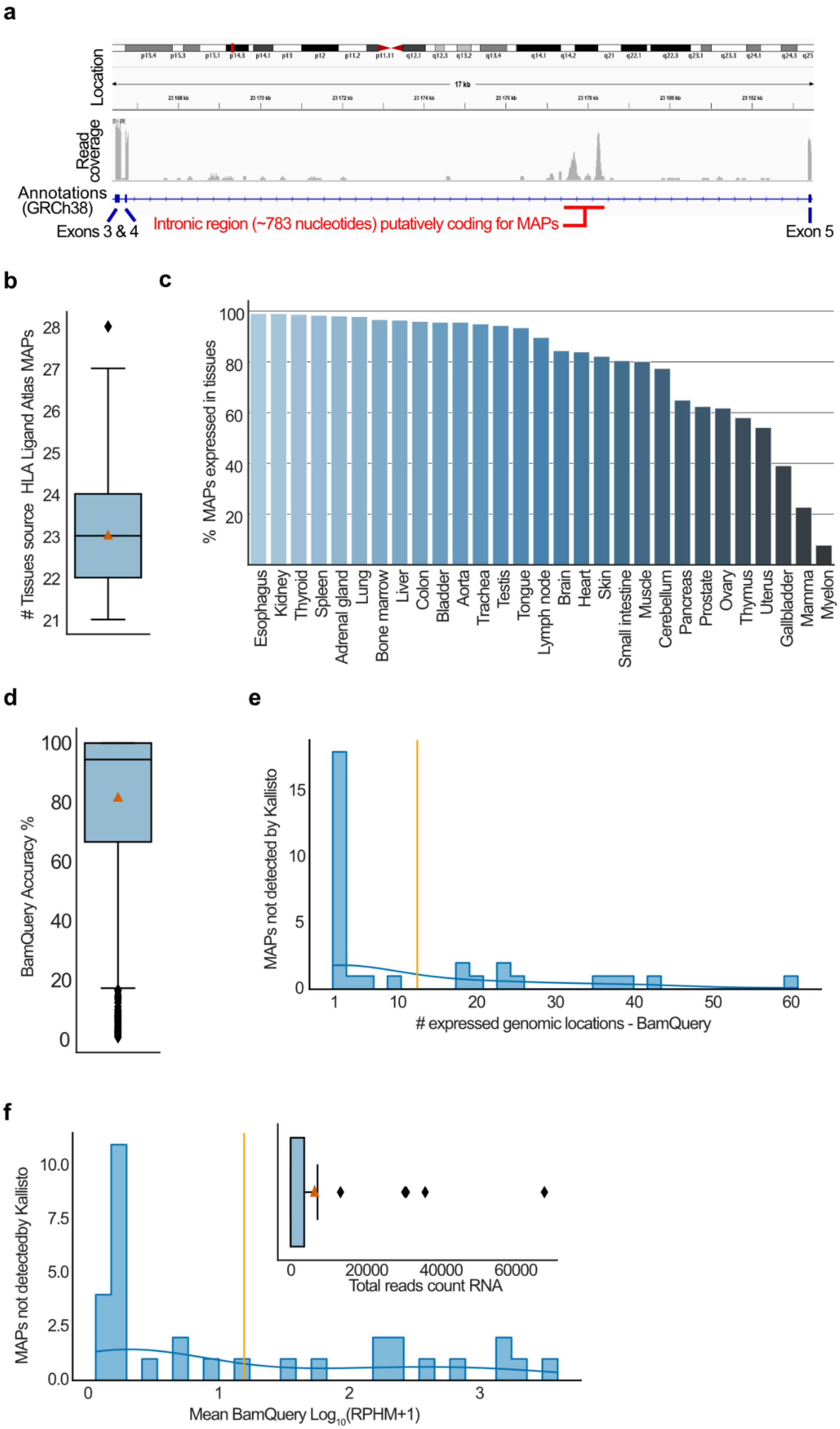

a
Location
17 kb
Read coverage
Annotations (GRCh38)
Exons 3 & 4
Intronic region (~783 nucleotides) putatively coding for MAPs
Exon 5
b
### Tissues source HLA Ligand Atlas MAPs
21
22
23
24
25
26
27
28
c
% MAPs expressed in tissues
20
40
60
80
100
Esophagus
Kidney
Thyroid
Spleen
Adrenal gland
Lung
Bone marrow
Liver
Colon
Bladder
Aorta
Trachea
Testis
Tongue
Lymph node
Brain
Heart
Skin
Small intestine
Muscle
Cerebellum
Pancreas
Prostate
Ovary
Thymus
Uterus
Gallbladder
Mamma
Myelon
d
BamQuery Accuracy %
0
20
40
60
80
100
e
MAPs not detected by Kallisto
0
5
10
15
1
10
20
30
40
50
60
### expressed genomic locations - BamQuery
f
MAPs not detectedby Kallisto
0.0
2.5
5.0
7.5
10.0
0
1
2
3
Mean BamQuery Log10(RPHM+1)
0
20000
40000
60000
Total reads count RNA

**Extended Data Fig. 1 | Origin canonical MAPs and BamQuery's quality control**

**d-f**, Published MAPs reported as canonical (n=1,702) were searched with BamQuery in mTEC bam files in stranded with genome version GRCh38.p13, gene set annotations release v38_104, dbSNP release 151, keeping variants alignments, and allowing higher levels of MCS alignments by STAR.

**a**, Genome browser (IGV) illustration for the gene LINC02718 (chr11:23,166,352-23,183,625) in a sample of acute myeloid leukemia (GSM4432540 on GEO) of the heterogeneity of read coverage observed in a typical intronic region (between exon 4 and 5). This would make the usage of genomic annotations irrelevant to quantify the expression of the small region putatively coding for MAPs as most of the annotated intron is not, or lowly, covered by reads (depth of coverage represented in grey).

**b**, Number of tissues at the origin of the canonical MAPs from the HLA ligand atlas shared in at least 20 tissues (n=1,702). Orange triangle represents the average (23).

**c**, Percentage of MAPs (n=1,702) presented by the indicated tissues.

**d**, Percentage accuracy measured between BamQuery-acquired read counts and Jellyfish's K-mer counts for nine-mer MAPs (n=1,211). Orange triangle represents the average percentage accuracy (82%).

**e**, Number of genomic locations detected by BamQuery for the 32 MAPs undetected in Kallisto's TPM quantification. Orange line represents the average number of genomic locations (11).

**f**, BamQuery-acquired RPHM expression for the 32 MAPs undetected in Kallisto's TPM quantification. Orange line represents RPHM expression average (1.1). Inside panel: total RNA-seq BamQuery-acquired reads for the 32 MAPs undetected by Kallisto. Orange triangle represents the average total RNA-seq BamQuery-acquired reads (n=6,474).

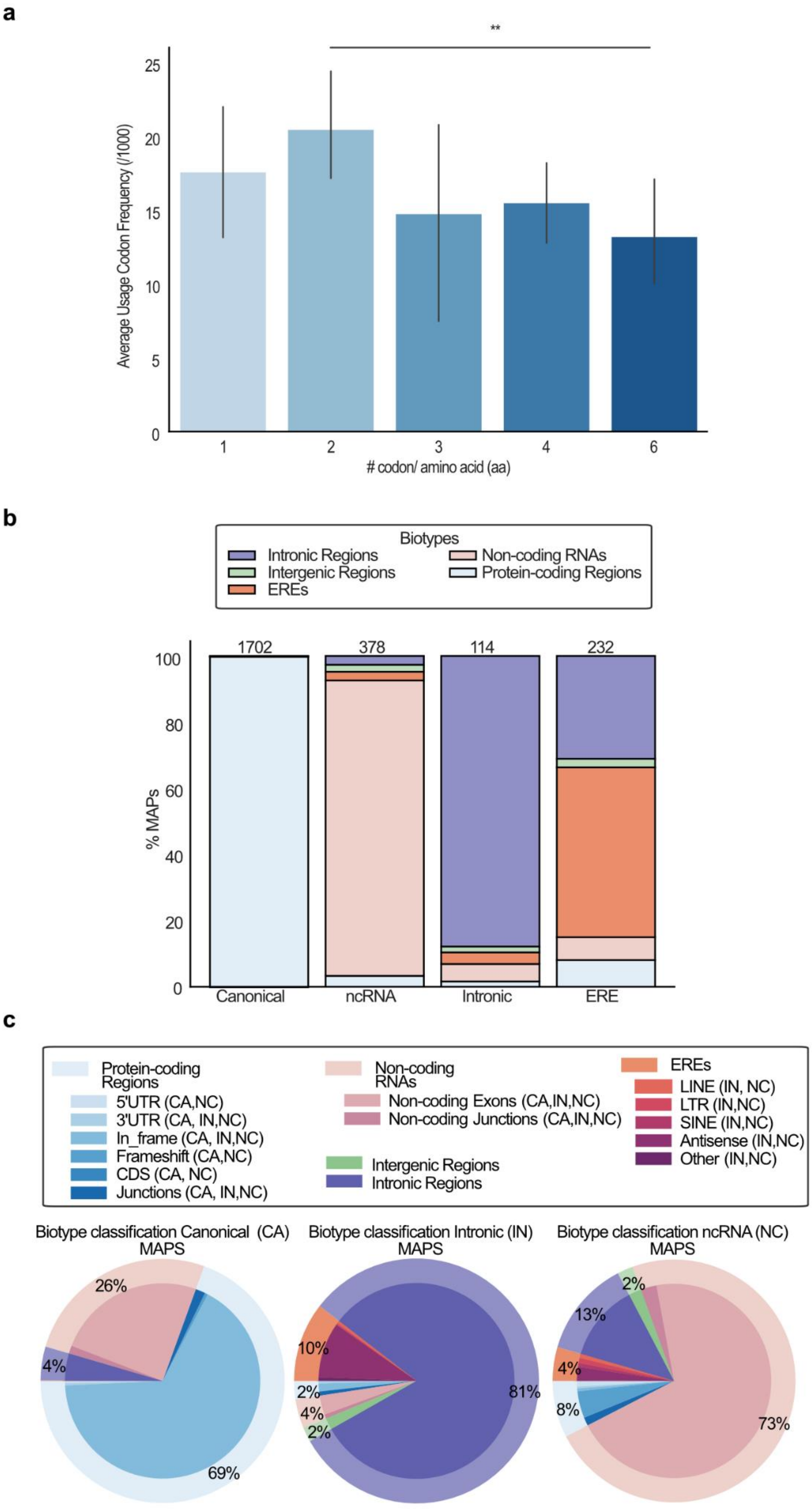

a
**
Average Usage Codon Frequency (/1000)
25
20
15
10
5
0
1
2
3
4
6
### codon/ amino acid (aa)
b
Biotypes
Intronic Regions
Intergenic Regions
EREs
Non-coding RNAs
Protein-coding Regions
% MAPs
100
80
60
40
20
0
1702
378
114
232
Canonical
ncRNA
Intronic
ERE
c
Protein-coding Regions
5'UTR (CA,NC)
3'UTR (CA, IN,NC)
In_frame (CA, IN,NC)
Frameshift (CA,NC)
CDS (CA, NC)
Junctions (CA, IN,NC)
Non-coding RNAs
Non-coding Exons (CA,IN,NC)
Non-coding Junctions (CA,IN,NC)
Intergenic Regions
Intronic Regions
EREs
LINE (IN, NC)
LTR (IN,NC)
SINE (IN,NC)
Antisense (IN,NC)
Other (IN,NC)
Biotype classification Canonical (CA) MAPS
26%
4%
69%
Biotype classification Intronic (IN) MAPS
10%
2%
4%
2%
81%
Biotype classification ncRNA (NC) MAPS
2%
13%
4%
8%
73%

**Extended Data Fig. 2 | Immunopeptidome properties of canonical and noncanonical MAPs.**

**a,** Average frequency of codons (among 1000 codons located in human reference protein-coding sequences) encoding each of the 20 amino acids. Codons of amino acids encoded by the same number of different synonymous codons were grouped together (x-axis).

**b**, Percentage of MAPs attributed to indicated biotypes by BamQuery based on the best guess biotype origin and on the genomic regions expressed in GTEx tissues and mTECs. The X-axis indicates the biotype reported in the original study (groups). For clarity, BamQuery biotypes were summarized into five general categories: protein-coding regions, non-coding RNAs, EREs, intronic and intergenic.

**c**, Percentage of the most likely biotype attributed by BamQuery to canonical, intronic and ncRNA MAPs.

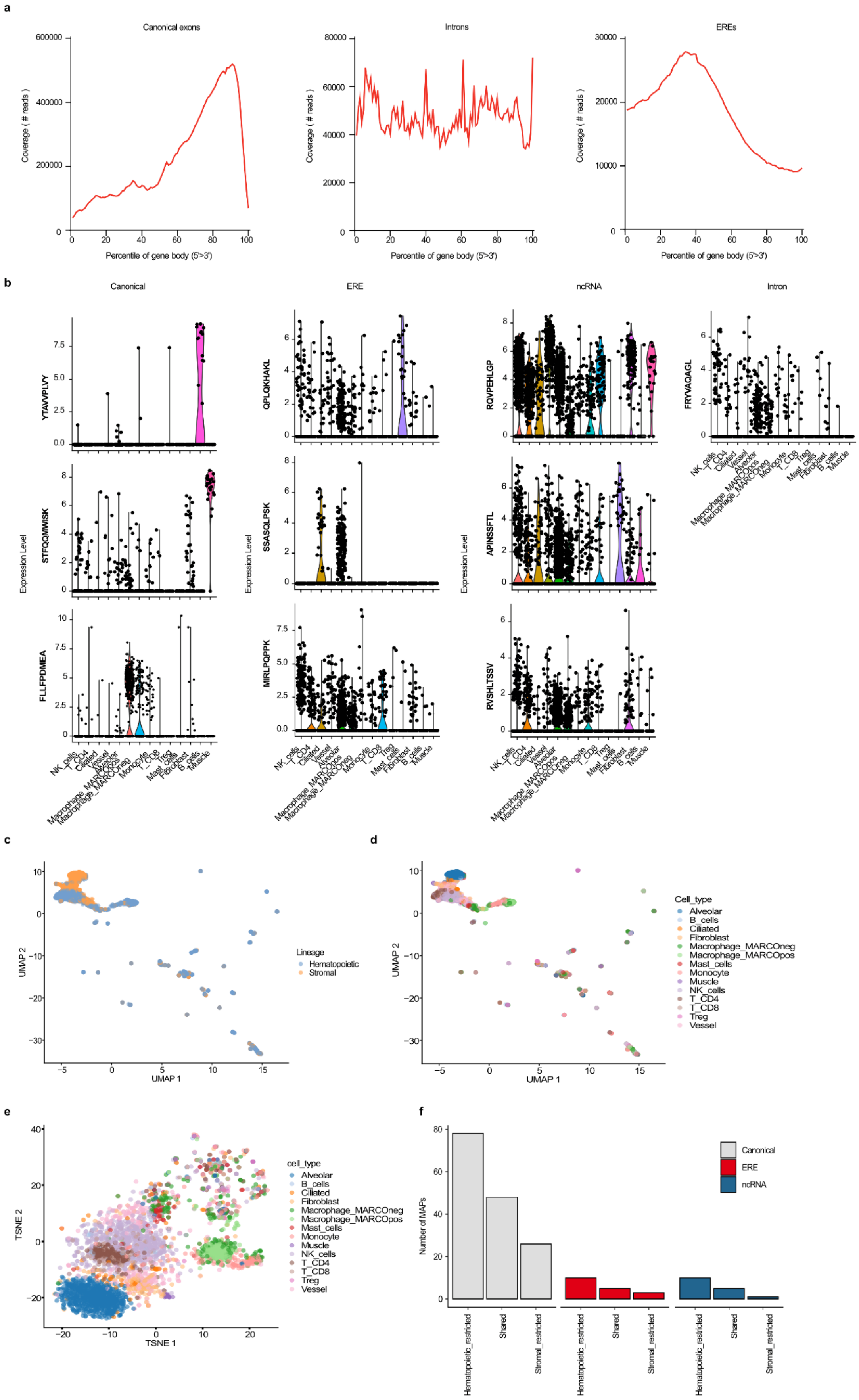
a
Canonical exons
Introns
EREs
Coverage ( # reads )
Percentile of gene body (5'->3')
b
Canonical
ERE
ncRNA
Intron
Expression Level
YTAVVPLVY
STFQQMWISK
FLLFPDMEA
QPLQKHAKL
SSASQLPSK
MIRLPQPPK
RQVPEHLGP
APINSSFTL
RVSHLTSSV
FRYVAQAGL
NK_cells
T_CD4
Ciliated
Vessel
Alveolar
Macrophage_MARCOpos
Macrophage_MARCOneg
Monocyte
T_CD8
Treg
Mast_cells
Fibroblast
B_cells
Muscle
c
d
Lineage
Hematopoietic
Stromal
UMAP 1
UMAP 2
Cell_type
Alveolar
B_cells
Ciliated
Fibroblast
Macrophage_MARCOneg
Macrophage_MARCOpos
Mast_cells
Monocyte
Muscle
NK_cells
T_CD4
T_CD8
Treg
Vessel
e
f
cell_type
TSNE 1
TSNE 2
Canonical
ERE
ncRNA
Number of MAPs
Hematopoietic_restricted
Shared
Stromal_restricted

**Extended Data Fig. 3 | BamQuery analysis of normal and cancer lung single-cell datasets.**

**a,** Number of lung scRNA-seq reads covering canonical genes, Intronic regions, and EREs.

**b**, Expression of canonical, ERE, ncRNA, or intronic MAPs identified as differentially expressed in the normal lung dataset.

**c**, UMAP depicting the clustering of the hematopoietic and stromal cells from the normal lung based on their MAP expression.

**d,** UMAP showing the clustering of the cell populations from the normal lung based on their MAP expression.

**e,** TSNE showing the clustering of the cell populations from the normal lung based on their MAP expression.

**f,** Number of canonical, ncRNA, or ERE MAPs identified by the differential expression analysis as restricted to the hematopoietic or stromal compartments or shared by cells of both lineages.

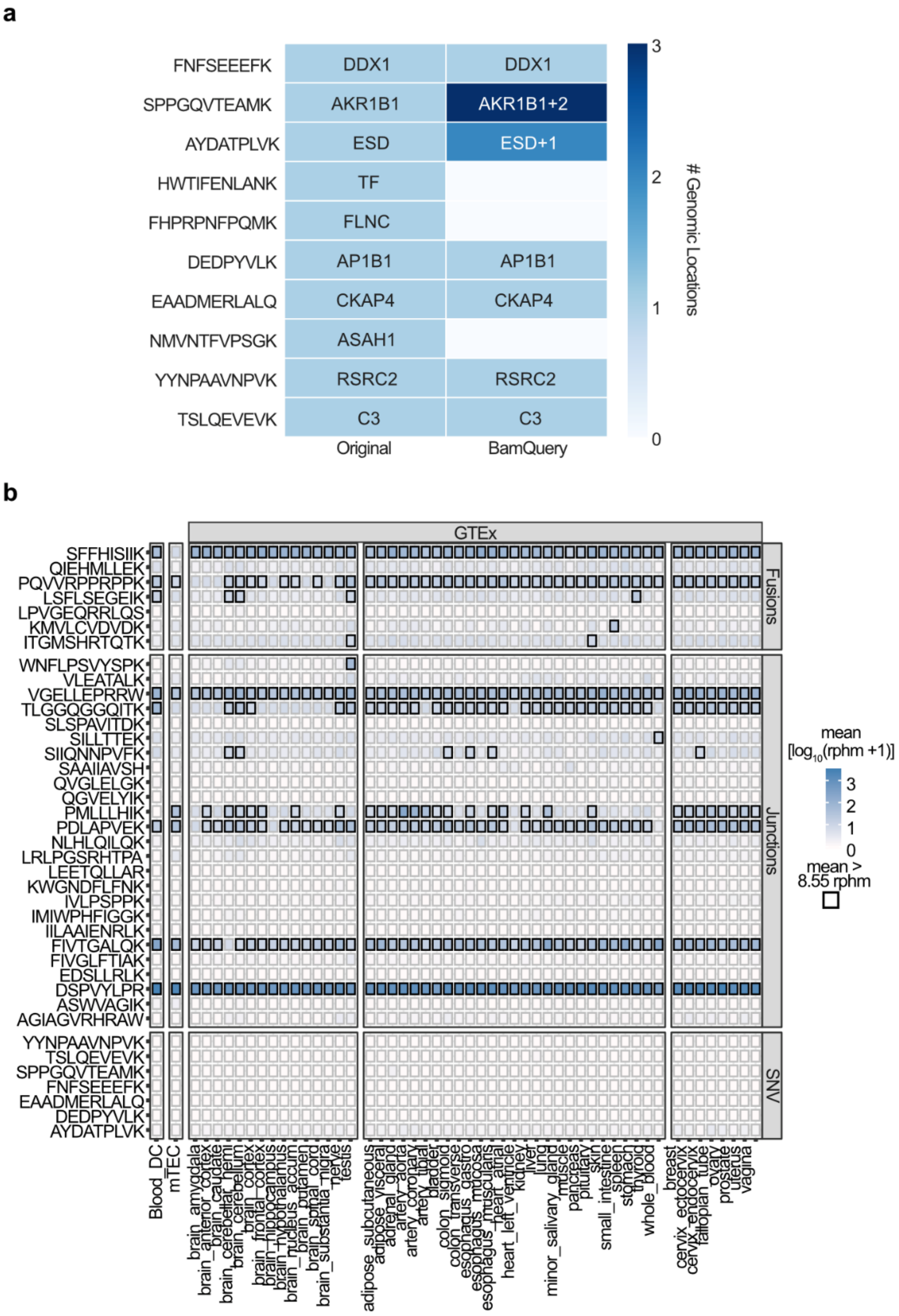


**Extended Data Fig. 4 | BamQuery elucidates safer immunotherapeutic targets.**

**a**, Heatmap of the number of genomic locations at which the expression of the SNVs-derived TAs was assessed by BamQuery vs by the original study.

**b**, Heatmap of average RNA expression of published fusions, junctions, and SNVs-derived TAs in indicated tissues. Boxes in which a peptide has an average rphm>8.55 are highlighted in black.

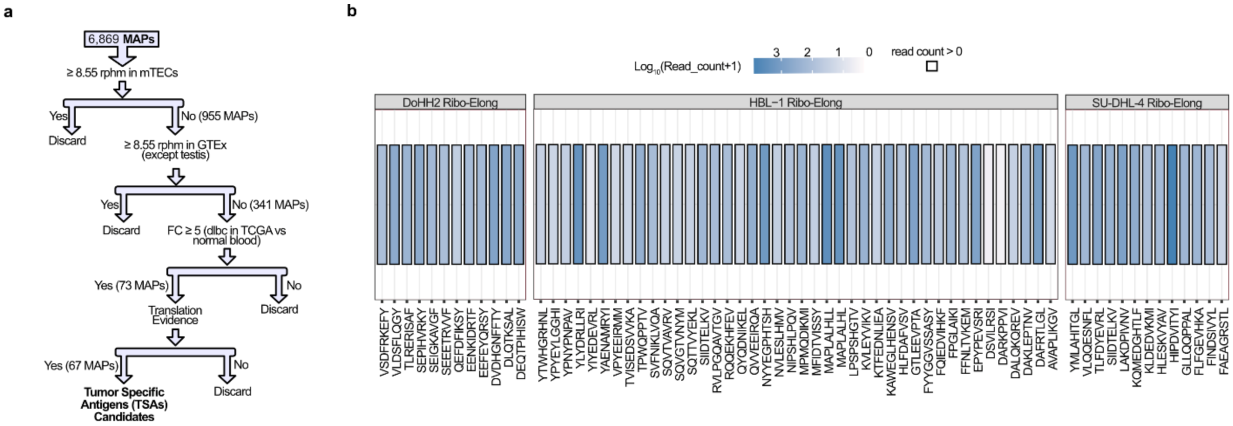


**Extended Data Fig. 5 | Discrimination of potential immunotherapeutic targets in DLBCL.**

**a**, Decision tree to discriminate TSAs from DLBCL.

**b,** Heatmap of average BamQuery-acquired read count of the 67 TSA candidates in indicated samples. Boxes in which a peptide has an rphm >8.55 are highlighted in black.

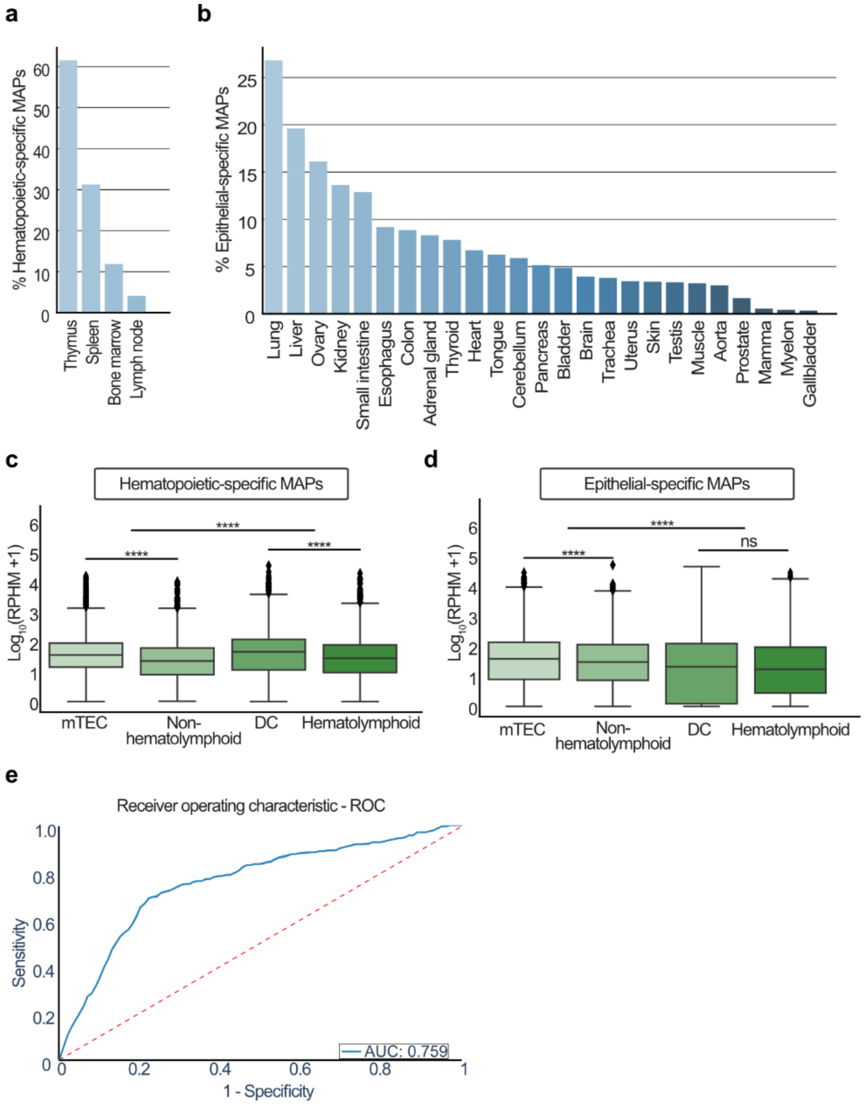


**Extended Data Fig. 6 | mTECs and Blood_DC for TAs prioritization.**

**a-b**, Percentage of hematopoietic-specific MAPs (n=2,429) (**a**) and epithelial-specific (n=3,237) (**d**) presented by the indicated tissues.

**c-d**, Average RNA expression of hematopoietic-specific (**c**) and epithelial-specific (**d**) MAPs in mTECs (n = 11), non-hematolymphoid GTEx tissues (n = 2,389), DCs (n =19) and hematolymphoid GTEx tissues (n=196). Wilcoxon rank-sum test two-sided was used for comparisons (****p<0.0001).

**e**, Receiver operating characteristic curve (ROC) for prediction of immunogenic based on RNA expression (RPHM) of mTEC and DC samples. AUC== ~0.75 with a 95% confidence interval (CI): 0.7588 - 0.7591.

**a**

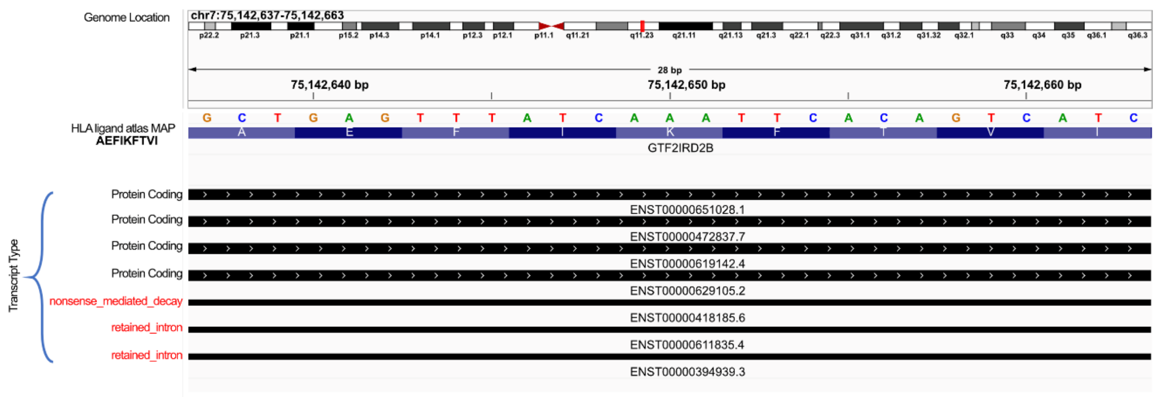


**Extended Data Fig. 7 | Different biotypes overlap at the same genomic location.**

**a**, The AEFIKFTVI peptide (HLA ligand atlas) at the indicated genomic location overlaps with protein-coding and non-coding RNAs transcripts.
