## Supplementary Tables Legends for "BamQuery: a proteogenomic tool for the genome-wide exploration of the immunopeptidome"

Supplementary Table 1. Origin of the MAP datasets used in the present study. 

Supplementary Table 2. Origin of the RNA-seq samples used in the present study. 

Supplementary Table 3. Enrichment of indicated MAPs in indicated clusters, computed from scRNA-seq data of normal lungs. 

Supplementary Table 4. Enrichment of indicated MAPs in indicated clusters, computed from scRNA-seq data of malignant lungs. 

Supplementary Table 5. Origin of the mutated TAs evaluated in the present study.

Supplementary Table 6. Report of the number of MAPs remaining after each filtering step in the TSA identification pipeline based on BamQuery. 

Supplementary Table 7. List of DLBCL TSAs identified through a BamQuery-based TSA identification pipeline. 

Supplementary Table 8. List of DLBCL CTAs identified through a BamQuery-based TSA identification pipeline. 

Supplementary Table 9. List of GO-terms enriched in the fraction of TCGA DLBCL patients expressing the highest levels of TSAs (top median) identified with BamQuery. 
